# Engineering microbial consortia for distributed signal processing

**DOI:** 10.1101/2025.04.23.650302

**Authors:** Katherine E. Duncker, Ashwini R. Shende, Irida Shyti, Ashley Ruan, Ryan D’Cunha, Helena R. Ma, Harshitha Venugopal Lavanya, Sizhe Liu, Neil Gottel, Deverick J. Anderson, Claudia K. Gunsch, Lingchong You

**Affiliations:** Department of Biomedical Engineering, Duke University, Durham, NC, USA; Center for Quantitative Biodesign, Duke University, Durham, NC, USA; Department of Computer Science, Duke University, Durham, NC, USA; Department of Mathematics, Duke University, Durham, NC, USA; Department of Civil and Environmental Engineering, Duke University, Durham, NC, USA; Division of Infectious Diseases, Department of Medicine, Duke University School of Medicine, Durham, NC, USA; Duke Center for Antimicrobial Stewardship and Infection Prevention, Duke University School of Medicine, Durham, NC, USA

## Abstract

A critical goal in biology is deducing input signals from measurable readouts. Genetic circuits have successfully been designed to respond to specific analytes and produce quantifiable outputs; however, multiplexed signal processing remains challenging. This limitation is partially due to crosstalk, or sensors’ non-specific responses to unintended signals. While strategies to achieve orthogonality are promising, they are time-intensive and context-dependent. Here, we introduce a new, generalizable approach that leverages microbial consortia to distribute sensory functions and computational methods to disentangle signals. Compartmentalizing sensor circuits within distinct populations simplifies experimental optimization by allowing individual populations to be exchanged without requiring genetic modifications. Our computational pipeline combines mechanistic modeling with machine learning to decode microbial communities’ unique temporal responses and predict multiple input concentrations. We demonstrated this platform’s versatility in a variety of contexts: measuring signals with high crosstalk, detecting antibiotics with natural microbial communities, and quantifying chemicals in hospital sink wastewater. Our findings highlight how combining microbial engineering with computational strategies can produce robust, scalable biosensors for diverse applications.

## Introduction

A fundamental challenge in biology is to deduce input signals from measurable outputs. The development of biological signal processing tools is essential for bioengineering research, medical diagnostics, and environmental monitoring^1–4^. Thus, a major effort in synthetic biology has been to engineer gene circuits that generate specific readouts in response to defined inputs, utilizing microbes, mammalian cells, and cell-free components^5–8^. As a result, dozens of sensory components have been constructed to detect diverse signals, including pH, gene transfer, heavy metals, explosives, and disease biomarkers^9–14^.

While tremendous progress has been made toward engineering biological sensing functions to detect individual analytes^15–17^, many applications require multiplexed signal processing, where multiple signals are measured simultaneously by a common system^18–20^. For example, diseases may be better characterized by a panel of analytes than by any individual signal alone^21^. Sensors have been created to monitor gut diseases by monitoring one analyte, yet the gastrointestinal environment is highly complex, with different niches containing a variety of nutrients, inflammatory markers, and microbial species^22–25^. Large gene circuits composed of logic gates have been programmed to process multiple input signals for cancer diagnostics^26^ and environmental contaminant detection^27^, although they were limited to classification tasks and did not yield quantitative measurements of each input.

Further progress in multiplexed signal processing has been limited due to persistent challenges related to crosstalk – a system’s non-specific response to unintended signals^28^. Multiplexed signal processing can be achieved by encoding information processing units in a single strain^18^ or distributing them across a microbial community^29–32^, but each approach has drawbacks. In single strains, each sensor circuit increases the growth burden imposed on cells, limiting the ability to scale the number of input signals processed^33,34^. Shared intracellular machinery and global factors like growth burden and metabolic competition also contribute to crosstalk between sensor circuits^35–40^. Crosstalk introduces bias into the system’s outputs, leading to incorrect inference of input signals. This interconnectedness also makes it challenging to optimize the performance of an individual sensor component without affecting the performance of other system components.

To this end, microbial communities reduce intracellular crosstalk and growth burden by compartmentalizing circuits, which also enables independent optimization of each circuit^30–32^. However, this approach introduces new challenges including intercellular interactions, like competition, that can lead to unstable community dynamics and impact the direct relationship between sensor inputs and outputs^27^. Output measurements distorted by imbalanced populations inhibit accurate resolution of inputs. This and other forms of crosstalk reduce measurement precision and thus limit the ability to quantify inputs on a continuous scale. Crosstalk can also be introduced by input-input interactions, environmental interference, output signal profile overlap, and other confounding factors. Thus, regardless of the method used, crosstalk is a universal challenge.

Efforts to reduce crosstalk to achieve orthogonality, where sensors operate independently, have focused on strategies to optimize circuits across all levels of cellular regulation, including modifying DNA sequences and tuning transcription and translation^41–45^. This typically requires time and resource intensive experiments to build and screen sensor components to minimize crosstalk^20^. Even when individual sensory components are engineered to minimize direct interactions with each other, orthogonality remains elusive due to global factors of the system and the vast potential for non-specific interactions^35–40^. Minimizing crosstalk is also context dependent: different sensor combinations, growth conditions, or measurement methods may require re-optimization^36–38^. Ultimately, if it is not possible to minimize crosstalk, then traditionally, that group of sensors would not be used together. Therefore, a more robust and efficient method is needed to overcome the challenges of crosstalk.

In this work, we develop a new strategy to address these challenges using computational signal resolution, which we demonstrate with perceptive microbial consortia. We asked whether a group of sensors with crosstalk could be recovered, not by experimental optimization of genetic components or populations, but through a computational approach that integrates mechanistic modeling and machine learning (ML). The only requirement for our approach is that a unique combination of inputs results in a unique combination of outputs. We illustrate the use of the approach on a series of increasingly complex experimental systems, highlighting the platform’s robustness and generalizability.

## Results

Our pipeline entails three steps (Fig. 1). First, we experimentally generated data by measuring cell density and fluorescence of a microbial community treated with different combinations of chemical input concentrations. Second, we augmented the experimental data with simulations using a mechanistic model. Third, we used a variational autoencoder (VAE) to encode the outputs of the augmented dataset into a compressed latent space, then trained a multi-layer perceptron (MLP) to use the latent variables to predict the chemical input concentrations. We demonstrated this workflow on systems with increasing complexity.

**Figure 1:**
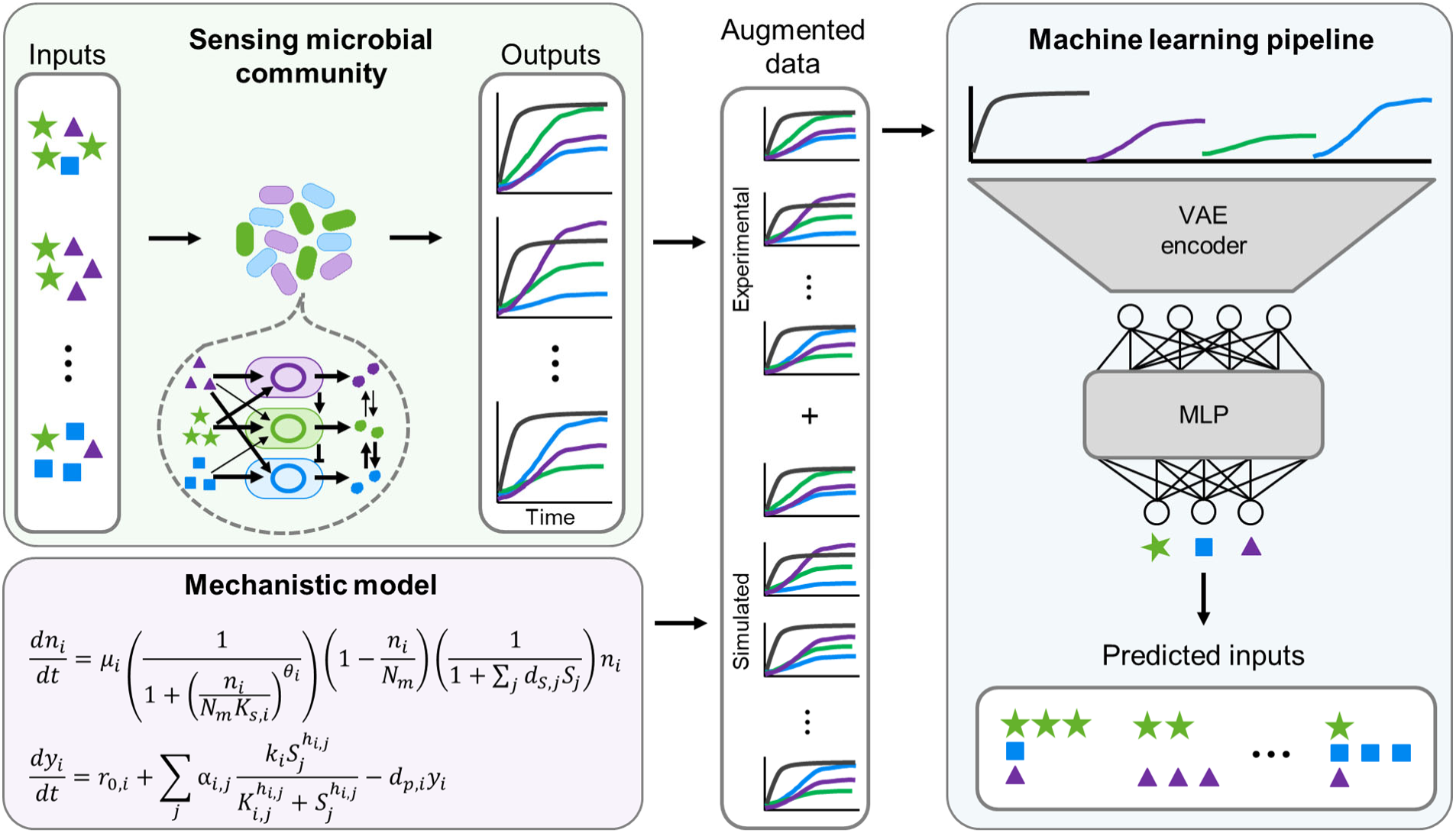
Overview of experimental and computational pipeline. (Top left) Microbial communities with sensing capabilities detect multiple inputs and produce outputs in response. These signal processing systems have natural complexities, including crosstalk between different sensors and signals. As long as each unique combination of inputs results in a unique output response, then it is mathematically possible to predict the inputs from the outputs. To maximize output uniqueness, we measured temporal responses. (Bottom left) To achieve sufficient data size to train our ML model, we used a mechanistic model which was fit to experimental data to augment the dataset with simulated data. (Right) The time courses are compressed into a lower dimensional latent space using a variational autoencoder (VAE). The latent dimensions are fed to a multi-layer perceptron to predict the experimental input concentrations.

### Pipeline development on a two-member orthogonal community

#### 1. System assembly and data generation

As proof of principle, we used a community of two *Escherichia coli* MC4100Z1 strains containing orthogonal inducible systems. One strain, aTc-GFP, expresses superfolder green fluorescent protein (sfGFP) under an anhydrotetracycline (aTc) inducible promoter; the other, IPTG-mCherry, expresses mCherry, a red fluorescent protein, under an isopropyl β-d-thiogalactopyranoside (IPTG) inducible promoter (Fig. 2a). These two inducible systems are commonly used, in part, due to their orthogonality. However, when we measured the community’s response to each inducer, it had a low level of crosstalk, where each fluorescent signal was negatively suppressed by the opposite inducer (Fig. 2b).

**Figure 2:**
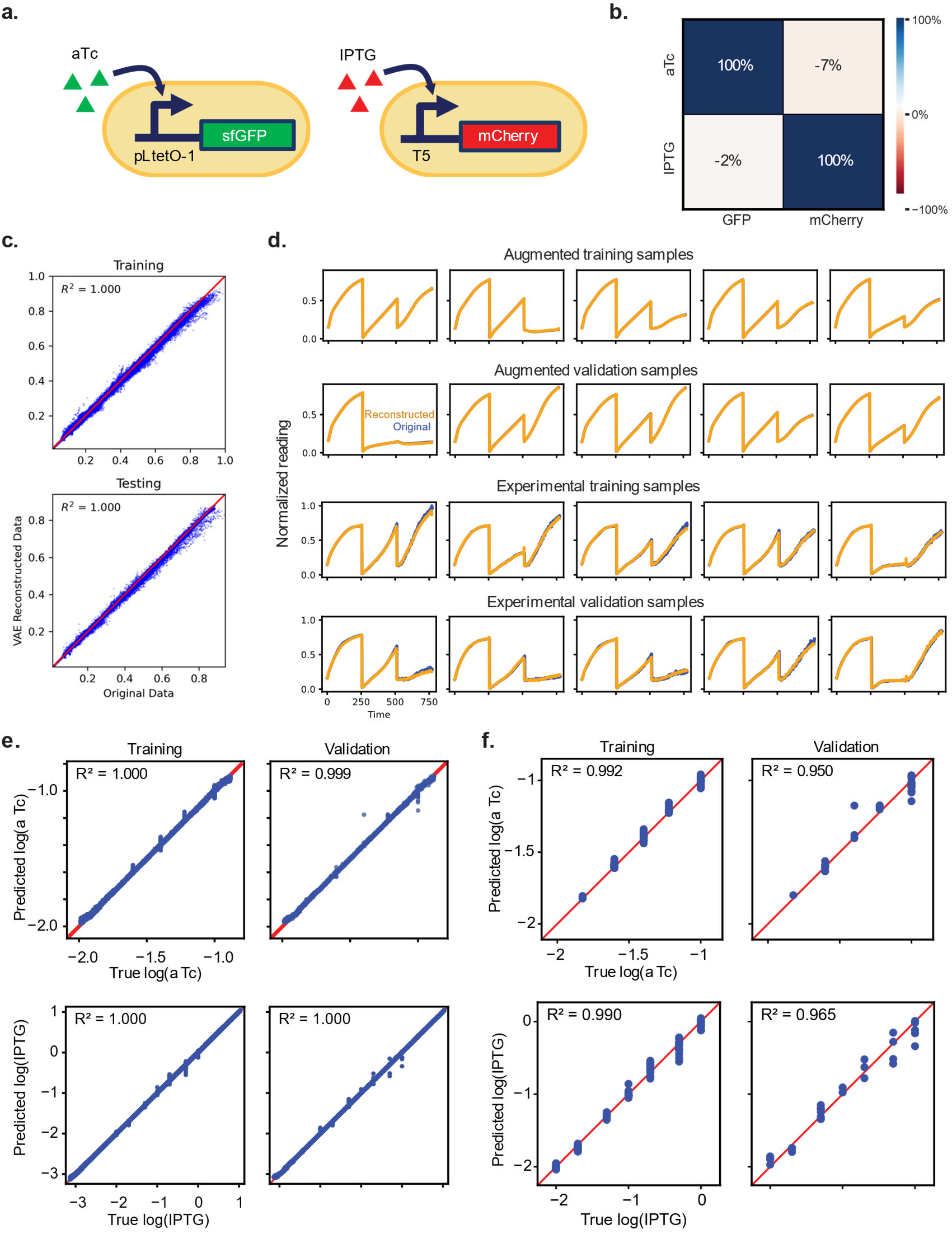
Pipeline can predict inputs from a 2-sensor community with low crosstalk. **a.** Schematic of low-crosstalk two-sensor community used to develop the experimental and computational pipeline. One sensor detects aTc and produces GFP in response, and the other sensor detects IPTG and produces mCherry in response. **b.** Crosstalk measurements for aTc and IPTG sensors. This set of sensors has minimal crosstalk. **c.** VAE reconstructed vs. true time course data with R^2^. Using the latent dimensions the VAE generated from time courses, the VAE decoder accurately reconstructs the features of the full time course data. **d.** Examples of VAE reconstructed time courses (orange) and their respective experimental time courses (blue). The VAE reconstructs the shape of the time courses with high fidelity. **e.** MLP predictions compared to true input concentrations of aTc and IPTG in the augmented dataset. Prediction accuracy was assessed with R^2^ metric. **f.** MLP predictions compared to true input concentrations of aTc and IPTG for experimental data only. The prediction accuracy achieved an R^2^ of at least 0.99 for both aTc and IPTG training data and 0.95 and 0.965 for aTc and IPTG concentrations, respectively, in the validation dataset.

Our approach to resolving crosstalk requires that a unique combination of inputs generates a unique combination of outputs. To achieve this, we designed experiments to maximize the information content of the outputs. In addition to using a different fluorescent reporter to respond to each input, which aligns with common practice, we also measured full time courses of cell density and fluorescence. Each time course contains far more information than a single time-point measurement^46^, which is the standard approach for estimating inputs. Moreover, time courses are more robust to experimental variabilities, such as choice of measurement time.

To generate data, we mixed cultures of the two populations at an equal ratio and added a range of concentrations of IPTG and aTc (Methods). In each experiment, we generated data for 95-380 samples, each consisting of the community growth curve, GFP, and mCherry time courses (Supplementary Figs. 1-3).

#### 2. Data augmentation using mechanistic modeling

While we were able to generate experimental data in high throughput, the VAE in our ML pipeline required significantly more data to be reliably trained. To address the limited experimental data, we augmented the dataset using a mechanistic model. The model consists of a set of ordinary differential equations (ODEs) to describe each population’s growth and protein expression (Methods).

The induction of each fluorescent protein is modeled using a Hill equation, constrained by measurements of the community dose-response at a single time point (Supplementary Fig. 4). Our model also accounts for crosstalk between the two channels, constrained by the measurements of the community’s fluorescence response to saturating concentrations of each inducer at a single time point (Fig. 2b). We used nonlinear optimization to estimate the remaining parameters of the model using the experimentally measured time courses. These parameters include growth rate, growth inhibition, basal protein expression, maximum protein induction, and protein decay rate.

The goal of parameter fitting is to ensure that our model is constrained by available experimental data. The Hill equation parameter estimations using the single time point dose response data first imposed a soft constraint on the model; then, the overall performance of the model was more strongly constrained by the nonlinear optimization of the full equations with the time courses. As a result, the fitted model allowed us to densely sample the dose responses of the system with high fidelity. Using the optimized parameters, we simulated 10,000 time courses with a range of randomized IPTG and aTc concentrations. The simulated and experimental time courses were combined to create an augmented dataset.

We used the optimized model to evaluate population-level growth dynamics of the microbial community. Although the measured crosstalk of the two populations was low, the model indicates that they could have underlying interactions, like imbalanced growth (Supplementary Fig. 5). Growth competition between populations is a challenge for implementing complex signal processing with microbial communities because it can cause one population to overtake the community^47–49^. This typically requires optimization at the community level to maintain stable population fractions^47,50^. Additionally, it may confound the relationship between inputs and outputs, thereby hindering accurate inference. However, we aimed for our computational pipeline to facilitate accurate signal processing despite such confounding factors present in the community.

#### 3. Using machine learning to predict signal concentrations from time courses

Our next objective was to use the augmented data to train a neural network to predict input signals directly from the system outputs. The outputs consisted of the community growth curve, GFP, and mCherry time courses. Each output was generated with high temporal resolution, consisting of at least 255 time points, to ensure rich information content. This resulted in high dimensional data, where the combined structure of the outputs was 3 × 255. In processing the data, the cell density and fluorescence time courses associated with each sample or simulation were concatenated into a single data series, yielding a vector of 765 time point features (Supplementary Fig. 6).

To facilitate input predictions from this high dimensional output data, we built a VAE to deduce a concise, latent representation of the time courses. Our recent studies have demonstrated that the latent dimensions generated by properly trained VAEs can improve downstream analysis by extracting key features of the data through an unbiased approach, minimizing the impactof experimental noise^51,52^. Upon successful training (Methods), each combined data series was compressed by the encoder of the VAE to a low-dimensional latent vector. The decoder of the VAE then reconstructed the original data series from the latent vector.

We systematically tested VAE architectures and latent space sizes to determine a configuration for reliable downstream prediction (Supplementary Note). We did so using this orthogonal sensor dataset and a dataset for two sensors with strong crosstalk, described in the next section. In this study, we selected a latent size of 10, which represents roughly 76-fold compression of the original data. Despite this drastic compression, each latent vector captures the essential features of the original time courses, as evidenced by the high-fidelity reconstruction of the raw data (Fig. 2c,d).

Given each latent vector’s ability to reconstruct the original data series with high fidelity (using the trained VAE decoder), the latent vector can serve as the surrogate of the raw data series. We next built a neural network combining the trained VAE encoder and an MLP to first map microbial community outputs into a latent vector and then map the latent vector to the corresponding input concentrations. When training this combined model, we froze the parameters of the VAE encoder and only optimized the parameters associated with the MLP (Methods). We chose to use a four-layer MLP to capture the complexity of the microbial community while maintaining simplicity to avoid overfitting.

The VAE-MLP accurately estimated IPTG and aTc concentrations for the augmented dataset (Fig. 2e). High accuracy was maintained when we isolated predictions solely for experimental data (Fig. 2f). The computational pipeline, including the VAE-MLP architecture, that was developed for this dataset was used for all subsequent analyses.

### Demonstration of the pipeline on two sensors with strong crosstalk

We next tested the pipeline on two sensors with high crosstalk that detect tetrathionate (TTR) and thiosulfate (THS)^22^. These two sulfur-related compounds are linked to gut inflammation^22,25,53^ as well as steel corrosion^54–56^ and toxicity^57,58^ in wastewater. The sensors, TTR-YFP and THS-CFP, were contained in *E. coli* MG1655 host strains for community measurements (Fig. 3a). We measured the community’s fluorescence activation in response to each of the inducers (Fig. 3b). CFP had 85% cross-reactivity to TTR, indicating strong crosstalk. YFP had 8% cross-reactivity to THS.

**Figure 3:**
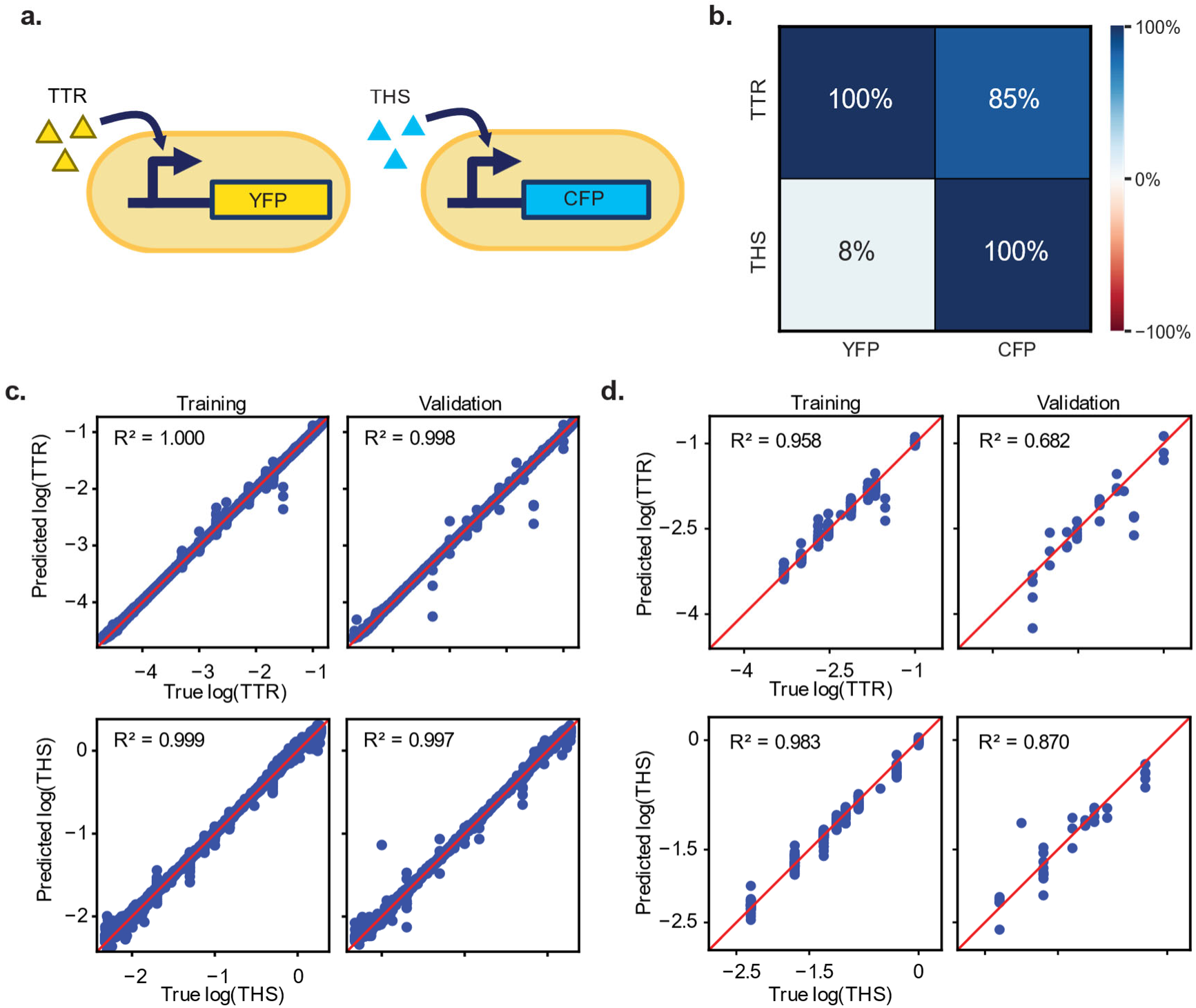
Pipeline can predict inputs for a 2-sensor community with high crosstalk. **a.** Two high-crosstalk sensor community to test the pipeline’s ability to handle crosstalk. One sensor detects TTR and produces YFP in response, and the other sensor detects THS and produces CFP in response. **b.** Crosstalk measurements for TTR and THS sensors. This set of sensors has high crosstalk as indicated by the strong activation of fluorescence by the unintended inducer. **c.** MLP predictions vs true input concentrations of TTR and THS for augmented training and test sets. Each have an R^2^ of at least 0.997. **d.** MLP predictions vs true input concentrations of TTR and THS for training and test sets of experimental data only. The model can achieve an R^2^ of at least 0.682.

We co-cultured the two sensor strains with a range of TTR and THS concentrations and measured the cell density, CFP, and YFP at 285 time points (Supplementary Figs. 7-9). The data was processed following the same computational pipeline described for the orthogonal sensors, including mechanistic model fitting, data augmentation, and VAE-MLP training and prediction. The full pipeline achieved an R^2^ over 0.99 for predicting TTR and THS concentrations in the augmented dataset (Fig. 3c). When isolating predictions for experimental data, it achieved an R^2^ of 0.96 and 0.98 for the training set of TTR and THS concentrations, respectively. It predicted TTR and THS concentrations in the validation set with an R^2^ of 0.68 and 0.87, respectively (Fig. 3d).

### Demonstration of the pipeline on three-sensor communities

We next applied the pipeline to communities consisting of three members, each containing one sensing circuit in *E. coli* Top10F’ host strains. To ensure the three reporters had distinct fluorescence wavelengths, we used red, cyan, and yellow fluorescent proteins for all three-sensor communities. We followed the same experimental protocol for generating time courses, except with three populations, three inducers, and measuring three fluorescence outputs in addition to cell density. Computationally, we adjusted the mechanistic model to incorporate the growth and protein expression of the third population. We used the same parameter fitting protocol with these updated equations and new datasets. The growth curve and three fluorescence curves were concatenated into a single data series of four total curves.

First, we constructed a 3-member community of sensors with low crosstalk (Fig. 4a). The strains, Cuma-YFP, OHC14-CFP, and aTc-mCherry, detect cuminic acid (Cuma), N-(3-Hydroxytetradecanoyl)-DL-homoserine lactone (OHC14), and aTc, respectively^18^. Our VAE-MLP pipeline enabled the prediction of the three input concentrations with high fidelity (Fig. 4b). Next, we combined three sensors with stronger crosstalk, up to 13%, between multiple fluorescence and inducer pairs (Fig. 4c). The strains, Van-YFP, DAPG-CFP, and Nar-mCherry, detect vanillic acid (Van), 2,4-diacetylphloroglucinol (DAPG), and naringenin (Nar), respectively^18^. Despite crosstalk, the computational pipeline accurately predicted the experimental input concentrations for these sensors (Fig. 4d).

**Figure 4:**
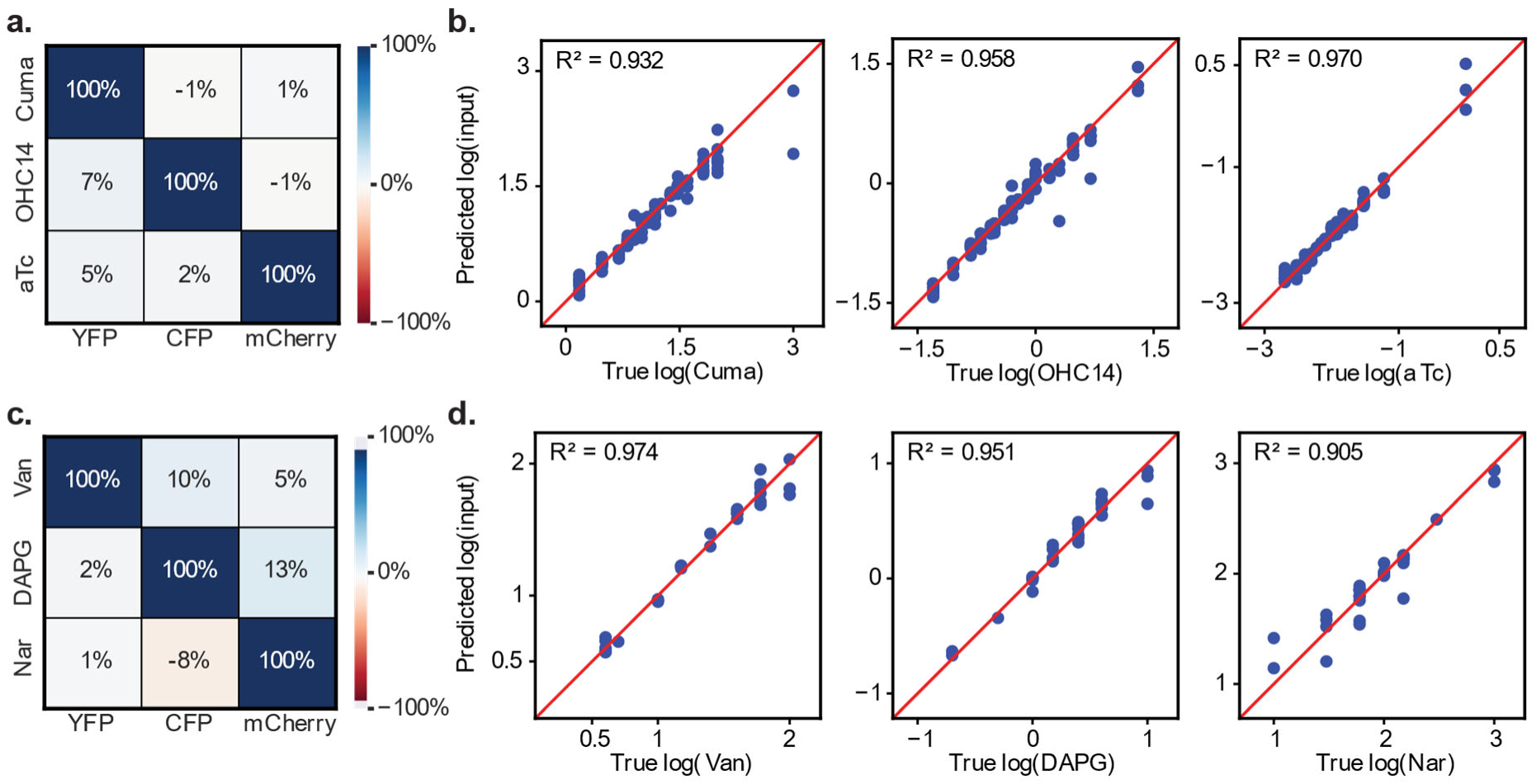
Input predictions for 3 sensor communities with varying degrees of crosstalk. **a.** Crosstalk measurements for three low-crosstalk sensors that detect Cuma, OHC14, and aTc. **b.** MLP predictions vs true input concentrations of Cuma, OHC 14, and aTc for the experimental test dataset. Each input type can be predicted with an R^2^ of at least 0.93**. c.** Crosstalk measurements for three high-crosstalk sensors that detect Van, DAPG, and Nar. **d.** MLP predictions compared to true input concentrations of Van, DAPG, and Nar for the experimental validation dataset. Each input type was predicted with an R^2^ of at least 0.90.

### Analyte quantification through indirect community responses

The fundamental requirement for our approach is a unique mapping between inputs and outputs, even without a direct sensing mechanism. To demonstrate this notion, we applied our pipeline to analyze a two-member community’s response to antibiotic combination treatment^59^. Neither population was engineered as a sensor for detecting chemical compounds. The community consisted of a β-lactam resistant population, expressing a β-lactamase (Bla), and a β-lactam susceptible population. The community was treated with varying concentrations of a β-lactam (amoxicillin) and a Bla inhibitor (tazobactam or sulbactam), which inhibits Bla expressed by the resistant population. Both populations constitutively expressed blue fluorescent protein (BFP), and the resistant population constitutively expressed GFP (Fig. 5a). Cell density, BFP, and GFP were measured for the duration of culturing.

**Figure 5:**
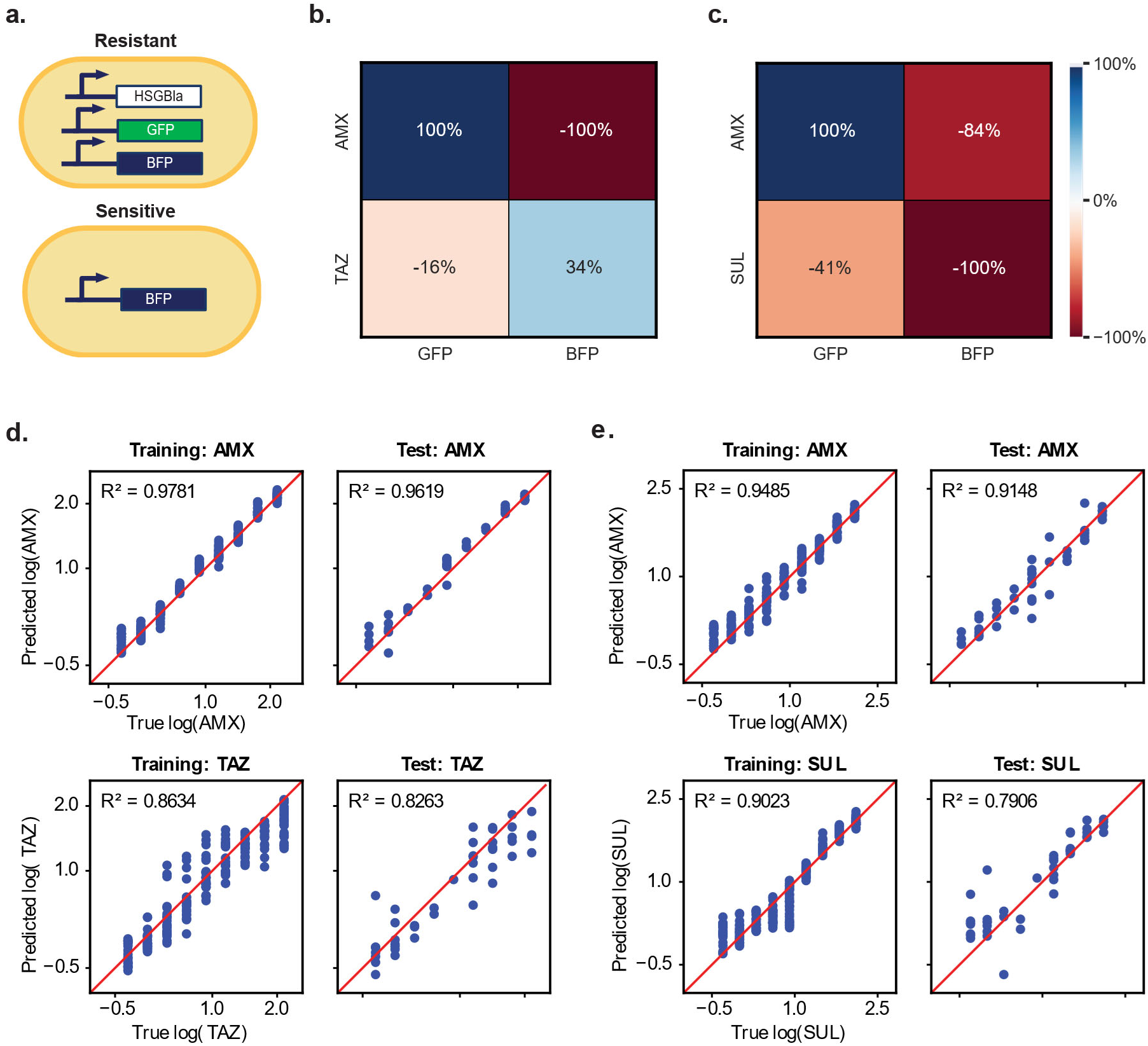
Pipeline predicts antibiotic and inhibitor concentrations from communities without direct sensing functions. **a.** The communities consisted of an antibiotic-resistant strain that expressed GFP and Bla, and an antibiotic-sensitive strain. Both strains expressed BFP. The resistant strain used for the results in this figure expressed a periplasmic Bla enzyme on a high copy number plasmid vector, named HSGBla. **b,c.** Heatmaps of the microbial community fluorescence responses to treatment with the maximum concentrations of the antibiotic or inhibitor (TAZ (**b**) and SUL (**c**)). Color bar and value represent the fluorescence fold-change with respect to no antibiotic or inhibitor added. Since the communities were not designed to detect the drugs, the same molecule could elicit the absolute maximum response for both fluorescent proteins (amoxicillin in **b**). The absolute maximum fold-change response could also be negative, indicating the molecule suppressed fluorescence (BFP maximum for both panels). **d.** MLP predictions of amoxicillin (AMX) and tazobactam (TAZ) Bla-inhibitor concentrations vs true concentrations for experimental training and testing datasets. **e.** MLP predictions of amoxicillin (AMX) and sulbactam (Sul) Bla-inhibitor concentrations vs true concentrations for experimental training and testing datasets.

We used our pipeline to predict the antibiotic and inhibitor concentrations from the three time courses. Although neither fluorescent protein was directly activated by the antibiotic or inducer, we estimated the crosstalk model parameter from community fluorescence data corresponding to the maximum antibiotic or inhibitor concentrations (Fig. 5b,c). We observed high levels of crosstalk in these communities, meaning they would not typically be selected as sensors for these two analytes. We also fit the Hill equation parameters to single time point fluorescence dose response data (Supplementary Fig. 10). Since the community was not designed to detect the antibiotic or inhibitor, the fluorescence responses to each molecule were noisier and less sensitive to lower concentrations. Our original two-population mechanistic model for sensing communities was sufficiently general to represent this new data, with different interpretations of variables (Supplementary Fig. 11). As such, we directly fit the remaining model parameters to this experimental data, then used the fitted model to augment the dose responses. Our VAE-MLP pipeline yielded high-accuracy predictions of the drug concentration combinations used to treat the communities (Fig. 5d,e).

We applied this approach to eight datasets of these two-member communities treated with amoxicillin and Bla inhibitors. In each dataset, the plasmid copy number, Bla enzyme, or inhibitor molecule were varied. In most cases, without modifying the mechanistic model formulation, our pipeline yielded accurate predictions of the drug concentrations, underscoring the generalizability and robustness of our approach (Supplementary Fig. 12). Certain plasmid and inhibitor combinations resulted in worse predictions, typically for the inhibitor. This is likely due to the inhibitor eliciting too small a change in cell density and fluorescence such that the pipeline cannot accurately estimate its concentration. For some datasets, the BFP measurements were noisier, preventing the pipeline from detecting differences in the signal response. Nevertheless, for most of the datasets, the concentrations were predicted with high fidelity.

In these examples, we show that general microbial communities with unique responses to inputs can be used with our pipeline to estimate important signals, such as antibiotic concentrations^60,61^. The communities do not need to be engineered with direct sensory circuits to act as signal processing systems. Additionally, using our default mechanistic model for fitting and data augmentation facilitates accurate predictions.

### Analyte quantification in hospital wastewater

Detecting analytes in complex samples like wastewater or patient samples is necessary for the translation of biological signal processing to environmental and biomedical applications. Real-world samples often contain many analytes at unknown concentrations, which typically interfere with the inference of input signals ^4,62–64^. Therefore, we sought to demonstrate that our platform can estimate analyte concentrations in complex real-world samples.

We tested this with water collected from sink P-traps in patient rooms from an acute care hospital. Hospital sink P-traps contain many contaminants, including pathogenic microbes, pharmaceutical compounds, and disinfecting agents^61,62,65^. Hospital sinks have been connected to antimicrobial-resistant outbreaks and are a known source for exposure to antibiotic resistance and spread of infections^66,67^. Also, hospital wastewater effluent is of concern due to its high concentrations of active pharmaceutical compounds that are toxic to wildlife and humans^61,68^. Quantifying analytes in hospital sink water could help toward studying the relationship between sinks and patient outcomes and toward mitigating against resistant outbreaks and toxic wastewater.

P-trap samples were collected weekly from newly installed sinks in patient rooms in Duke University Hospital (Methods). To test our platform, we pooled together the filtrate of five sinks from one week (Fig. 6a). We used the Van-YFP and aTc-mCherry sensors previously used in three-sensor community experiments. Vanillic acid (Van) is a phenolic compound which could be present from pharmaceuticals in hospital wastewater^69^, and aTc is a degradation product of tetracycline antibiotics^70,71^.

**Figure 6:**
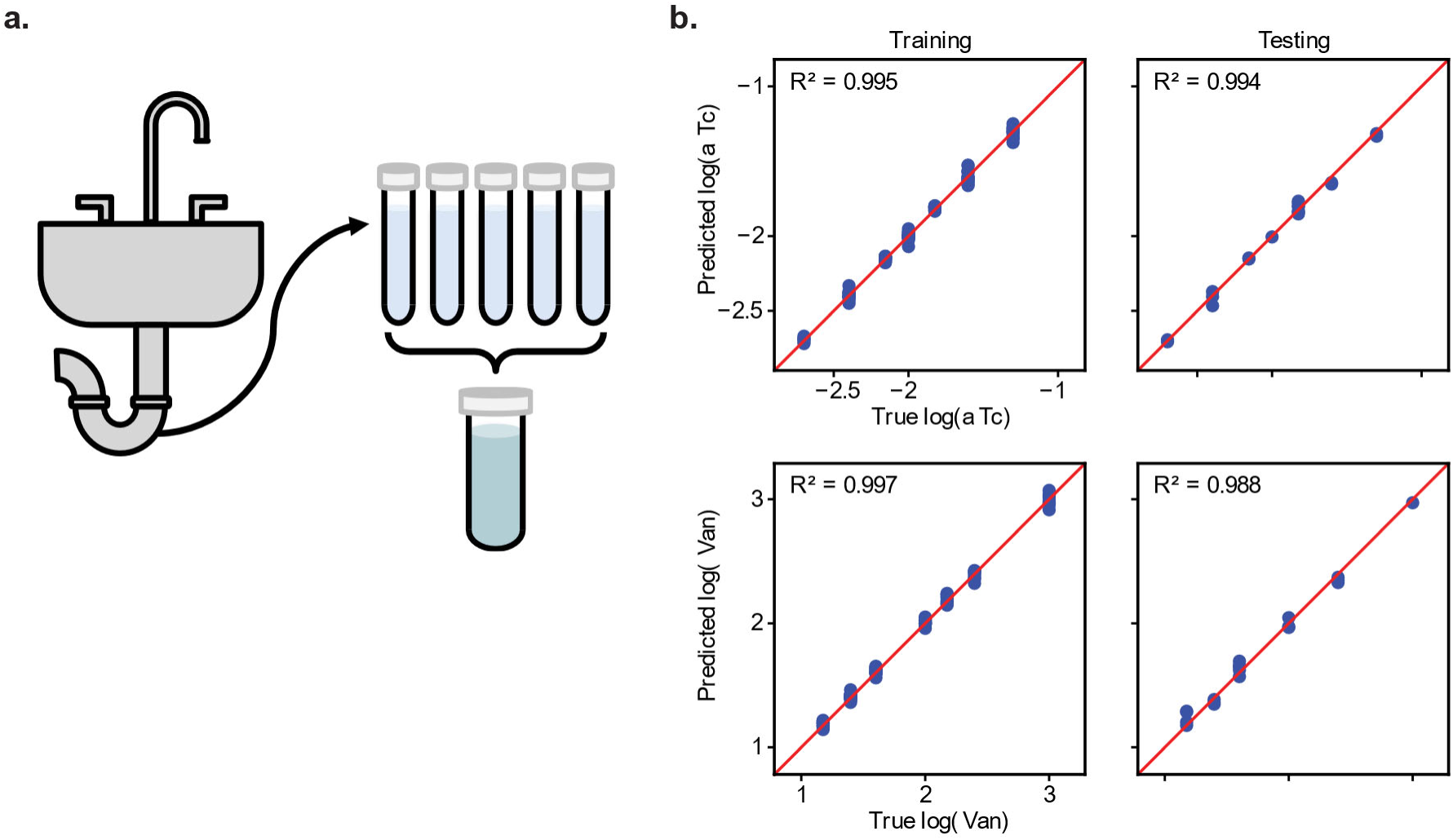
Platform can estimate two analyte concentrations in a complex sample collected from hospital sink P-traps. **a.** Schematic overview describing the sample collection and processing. Liquid was collected from the P-trap of hospital sinks. The samples from 5 sinks not treated with Virasept disinfectant were pooled together for testing. **b.** Final predictions of aTc and Van concentrations with respect to their true concentrations for both training and validation sets of the data. R^2^ score for each subset is labeled in the plot.

We first measured the sensors’ responses to Van and aTc in sink water (Supplementary Fig. 13). We added both sensor strains and various concentrations of Van and aTc to the pooled sink samples. We measured cell density and fluorescence time series to train and validate the computational pipeline. The pipeline predicted Van and aTc concentrations in sink water with an R^2^ of at least 0.99 for training data. For validation data, the pipeline estimated aTc concentrations with an R^2^ of 0.96 and Van concentrations with an R^2^ of 0.98 (Fig. 6b). This demonstrates that our pipeline is robust to measuring analytes in real-world samples with complex background compositions.

## Discussion

Our platform relies on several key design features critical to achieving multiplexed biological signal processing. By distributing sensory functions across microbial consortia, we reduced intracellular interactions between circuits, simplifying optimization, and eliminated the limit on scaling inputs imposed by growth burden. Augmenting experimental data using a mechanistic model generated datasets large enough to train a VAE. Our model describing microbial community growth and sensor activation sufficiently captured experimental data while maintaining simplicity to keep parameter fitting reliable. However, other models could be implemented to apply the pipeline to systems like cell-free genetic circuits or human cell-signaling pathways^5,8^. Finally, the VAE enabled distillation of important information from high dimensional data, and the MLP was critical for computing input concentrations from latent variables.

We used ML rather than directly using the mechanistic model to estimate input concentrations because of its flexibility, which enabled more accurate predictions (Supplementary Fig. 14). The VAE-MLP can better predict inputs because it is trained on the full training dataset, isn’t constrained by our mechanistic interpretation, and has more flexibility in learning correlations between inputs and patterns in the time courses. The mechanistic model only relies on the time series of the respective sample to estimate input concentrations for that sample, so it cannot infer generalized rules, and it is constrained by the equations we constructed. To make the pipeline more efficient, a pre-trained foundation model for encoding time series data^72–74^ could be used to skip the data augmentation and VAE training steps for each dataset. MLP training requires less data than VAE training, so latent variables encoded by a foundation model from experimental time courses could be directly passed through the MLP without augmentation.

Our pipeline overcomes many challenges in biological signal processing that could enable more generalizable translation of responsive biological systems to applications in diagnostics and environmental monitoring. We demonstrated how our pipeline could reduce bottlenecks that exist at various stages of traditional biosensor development. First, the platform expands the pool of candidates that can be considered as sensors because precise mechanistic understanding of specific genetic pathways or sensory components is not required. Biological systems with a natural response to stimuli can be implemented for detecting signals, even without a direct input to output sensing mechanism, as demonstrated with the antibiotic treated communities. Next, using this platform, sensors that have crosstalk with each other can be used in combination, whereas they otherwise would be excluded or require re-optimization. This broadens the sensor combinations available, enables scaling of the number of inputs measured, and minimizes the experimental optimization necessary. Finally, our concept increases the application opportunities for biological signal processing. Unknown interactions with other analytes in complex samples, which would interfere with input estimation, are resolved by our computational pipeline, as demonstrated on the sink water samples.

The concept we presented is agnostic of the type of inputs and outputs. We estimated small molecule concentrations, but the pipeline could be used as a general measurement tool in other contexts, such as estimating genetic diversity, gene expression, or pathogen abundance^19,75,76^. Alternative reporters with high information content, including hyperspectral profiles^77^, acoustic signals^78^, or high resolution fluorescence wavelength measurements, could be used instead of fluorescence time course data. Our platform could be implemented in many future directions as long as they meet the fundamental requirement that there is unique mapping between the measurable outputs and the inputs of interest.

## Methods

### Strains and plasmids

The IPTG-mCherry sensor uses p15a-T5-2A-mCherry plasmid developed in a previous study, in which it was referred to as His-T_2_-mCherry^79^. The aTc-GFP sensor uses p15a-pLTet-O-1-sfGFP plasmid, which we constructed from p15A-pTet-sfGFP-linker-Tdimer-kanR^80^ by removing the linker-Tdimer sequence. Both plasmids were kanamycin resistant and were transformed into *E. coli* MC4100Z1 host strains.

The THS-CFP sensor was constructed from pKD236-4b and pKD237-3a-2 plasmids; The TTR-YFP sensor was constructed from pKD238-1a and pKD239-1g-2 plasmids^22^. These were gifts from Jeffrey Tabor (Addgene references in Supplementary Table 1). The regulator plasmids, pKD237-3a-2 and pKD239-1g-2, were modified by replacing sfGFP with CFP and YFP, respectively, and the constitutive mCherry genes were removed. The plasmid pairs corresponding to each two-component system were chloramphenicol and spectinomycin resistant and were co-transformed into *E. coli* MG1655 host strains.

The OHC14-CFP sensor uses plasmid pAJM.1642; the Cuma-YFP sensor uses plasmid pAJM.657; the aTc-mCherry sensor uses plasmid pAJM.011; the Van-YFP sensor uses plasmid pAJM.773; the DAPG-CFP sensor uses plasmid pAJM.847; the Nar-mCherry sensor uses plasmid pAJM.661^18^. These were gifts from Christopher Voigt (Addgene references in Supplementary Table 1). The YFP reporter in pAJM.011 and pAJM.661 was replaced with mCherry to obtain aTc-mCherry and Nar-mCherry sensors. The YFP reporter in pAJM.1642 and pAJM.847 was replaced with CFP to obtain OHC14-CFP and DAPG-CFP sensors. The plasmids were all kanamycin resistant and were transformed into *E. coli* Top10F’ host.

Plasmid modifications were performed using Gibson Assembly cloning method^81^. Transformations were plated on LB agar plates with appropriate antibiotic selection. Colonies from the transformations were cultured in LB media overnight with antibiotic selection, then stored in 25% glycerol at −80°C.

Plasmids used in the antibiotic prediction dataset were originally generated and described in Ma *et al.*^59^. Our study only used data from *E. coli* MG1655 DA28102 cells with plasmids pBla-sfGFP (low copy number, expresses periplasmic Bla), pBlaM-sfGFP (low copy number, expresses cytoplasmic Bla), pHSGBla-sfGFP (high copy number, expresses periplasmic Bla), or pHSGBlaM-sfGFP (high copy number, expresses cytoplasmic Bla). DA28102 constitutively expresses mTagBFP2 on the chromosome, and all the plasmids express Bla under IPTG induction and constitutively express sfGFP.

### Growth media and chemical inducers

Unless stated otherwise, all agar plates were made with LB agar and all liquid cultures were in LB broth, both supplemented with appropriate antibiotics (50 µg/mL kanamycin, 35 µg/mL chloramphenicol, 50 µg/mL spectinomycin). Cultures for the sensor community experiments were induced with the following chemicals: isopropyl-β-D-thiogalactopyranoside (IPTG) (Thermo Scientific, Sigma, and GoldBio), anhydrotetracycline hydrochloride (aTc) (Sigma, 37919-100MG-R), sodium thiosulfate pentahydrate (THS) (Sigma, S6672-500G), potassium tetrathionate (TTR) (Sigma, P2926-25G), *N*-(3-Hydroxytetradecanoyl)-DL-homoserine lactone (OHC14) (Sigma, 51481-10MG), 4-Isopropylbenzoic acid (Cuma) (Sigma, 268402-5G), vanillic acid (Van) (Sigma, 94770-10G), 2,4-diacetylphloroglucinol (DAPG) (Santa Cruz Biotechnology, CAS 2161-86-6, Cat# sc-206518D), and naringenin (Nar) (Sigma, N5893-1G). The antibiotic prediction dataset used amoxicillin (Sigma) as the antibiotic and either tazobactam (Fisher Scientific) or sulbactam (Fisher Scientific) as the Bla inhibitors, which were prepared fresh in DMSO (amoxicillin, tazobactam) or water (sulbactam) for each experiment^51,59^.

### Sensor community experiments

Freezer stocks were streaked onto LB agar plates with antibiotic selection and incubated at 37°C overnight. Cultures were inoculated from single colonies and cultured overnight in 3 mL LB media with appropriate antibiotics in 16 mL culture tubes (Genesee Scientific, 21-129) at 37°C with 225 rpm shaking. Cultures were diluted with LB to reach OD600 (OD) of about 0.1. Appropriate antibiotics were added. For microbial community experiments, each population’s diluted culture was mixed at an equal volume ratio. The mixed cultures were transferred to a 96-well black-sided, clear flat bottom plate (Corning, 3603). Inducers were added to the appropriate wells to reach their intended final concentration (available in data files). Total volume for all microwell-plate cultures was 200 µL. 50 µL mineral oil was added on top of the cultures to prevent evaporation. Plates were incubated for at least 20 h at 37°C in a plate reader (Tecan Infinite 200 PRO), which shook the plate (5s, orbital) and measured OD and fluorescence every 5 minutes. Some experiments utilized Tecan Freedom EVO robot for liquid handling steps or for moving plates from incubator to plate reader for each measurement over the course of the experiment. When using the robot, plate readings were taken roughly every 17 minutes.

### Antibiotic treatment experiments

All data for the antibiotic treated communities was originally generated and described in Ma *et al.* and Baig *et. al*^51,59^. The methods used for the data included in our study are as follows. Colonies on LB agar plates were inoculated in 2 mL LB in 16 mL culture tubes for overnight culturing at 37°C with 225 rpm shaking for 16 h. 1 mM IPTG and 50 µg/mL kanamycin were added to cultures of the plasmid-containing population. Cultures were corrected to OD of 1. The resistant and susceptible population cultures were mixed 1:1, then diluted 16-fold for a cell density of 5 × 10^7^ cells/mL. 50 µL 1 M IPTG was added. Amoxicillin and Bla inhibitor stocks were diluted in LB at concentrations 2.5 times their final concentrations. 40 µL of the 2.5x stocks of amoxicillin and inhibitor were distributed to wells in a 384-well deep well plate (Thermo Scientific) using a MANTIS liquid handler (Formulatrix). Then, 20 µL of the diluted culture was added by the MANTIS. Initial cell density was 1 × 10^6^ cells per 100 µL well, IPTG concentration was 1 mM, antibiotic and inhibitor concentrations formed a dose-response matrix of 0, 0.5, 1, 2, 4, 8, 16, 32, 64, and 128 µg/mL each. Each condition had three replicate wells and were randomly positioned across the plate. All edge wells and 12 interior wells contained blank LB. The plate was incubated at 30°C in a plate reader (Tecan Spark) with the lid on to prevent evaporation. The plate reader, which was equipped with a lid lifter, shook the plate (5s orbital) and took OD, GFP, and BFP readings every 10 min for 24 h.

### Hospital sink water collection and processing

Sink samples were collected from the 9100 unit in Duke North tower of Duke Hospital in Durham, NC. 30 in-room patient handwashing sinks that did not have multi-drug-resistant organism colonization were used in the study and randomized 1:1 to intervention or control groups. Intervention sinks were treated with Virasept (Ecolab) disinfectant while control sinks received no treatment. Sink P-traps were sampled weekly using sterile syringes and tubing to collect the sample. The sample was vortexed in the syringe for 10 seconds then 15 mL was dispensed into a sterile 15 mL tube, which was vortexed again for 10 seconds, then stored at 4°C. Samples were filtered through a 0.22 µm paper filter into a flask with vacuum pressure applied. The filtrate from each sink sample was poured into a 15mL conical tube and stored at 4°C again until use for the experiments in this study.

Immediately prior to using the samples for the sensor experiments, they were filtered again through a 0.22 µm syringe filter 2x to ensure removal of any native sink microbes or contamination from processing that could interfere with the growth of the sensor bacteria. The filtrate samples from 5 control sinks (no treatment) from week 12 were mixed at an equal volume ratio to create an average hospital sink sample. The sink mixture was mixed with LB at a 3:1 ratio, to which appropriate antibiotics were added, then 176 µL was distributed to wells of a 96-well plate. Sensor bacteria were prepared as described for sensor community experiments, except cultures were corrected to reach ∼0.7 OD before mixing together at an equal volume ratio. 20 µL of the mixed cultures was added to a 96-well plate. 2 µL of each inducer was added to the corresponding wells. Mineral oil was added, and the plate was incubated and readings were taken in a plate reader over time.

### Experimental data pre-processing

Data was imported to Python for the remaining methods. For co-culture data, all time courses from a single set of sensors were trimmed to match the shortest time span from the sensor set and subsampled to have the same sampling interval. Fluorescence data obtained from different plate readers was converted to the same scale using a calibration between instruments (available in data files). Converted fluorescence data was min-max scaled with respect to the rest of the corresponding fluorescence data in the sensor set by performing the following scaling: 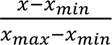. The basal expression fluorescence time course (when no inducers were added) was subtracted from all time courses. The OD and fluorescence time courses were concatenated to create a 1-dimensional vector per sample that is of length 𝑡x (𝑁 + 1), where 𝑁 is the number of fluorescence colors in a community and 𝑡 is the number of time point measurements.

### Mechanistic model formulation

We formulated a set of ordinary differential equations (ODEs) to describe each population’s cell density (𝑛_i_) and fluorescent protein per cell (𝑦_i_) in response to different input concentrations (𝑆_j_), where 𝑖 is the population and 𝑗 is the input:

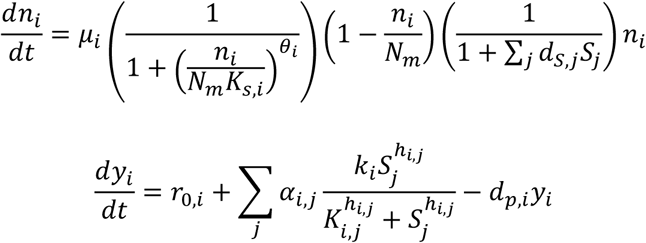

The cell density equation incorporates maximum growth rate (𝜇_i_), carrying capacity (𝑁_m_), and input-derived growth inhibition (𝑑_S,j_). A term with the parameters 𝐾_s,i_ and 𝜃_i_ is incorporated, which modifies the logistic growth equation to better fit the shape of experimentally measured OD time courses that we observed^51^. Fluorescence equations include basal gene expression rate (𝑟_O,i_) and protein decay rate (𝑑_p,i_). Fluorescence activation by each input is represented by a Hill equation with maximum induction rate (𝑘_i_), Hill coefficient (ℎ_i,j_) and the input concentration resulting in half-maximum induction (𝐾_i,j_). A crosstalk parameter (𝛼_i,j_) modulates how strongly each input activates expression with respect to the cognate input, where cognate inputs have an 𝛼_i,j_ of 1 and 𝛼_i,j_for non-cognate inputs is a relative fraction. To approximate measured optical density (OD) and total fluorescence (𝑌_i_), we use the following equations:

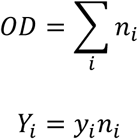

### Parameter estimations

Crosstalk, 𝛼_i,j_, was estimated from the community’s fluorescence measurements at 20 h for conditions where only one inducer was added at its maximum concentration. The average fluorescence value of the blanks was subtracted from the fluorescence values of the samples to get 𝑓𝑝(𝑆_j_). Crosstalk was calculated via the following equation,

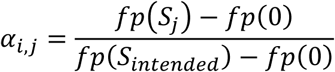

where 𝑓𝑝(𝑆_j_) is the fluorescence when the inducer of interest was added, 𝑓𝑝(0) is the fluorescence when no inducers were added, and 𝑓𝑝(𝑆_intended_) is the fluorescence when the inducer intended for that respective sensor was added.

The Hill coefficient (ℎ_i,j_) and half-max (𝐾_i,j_) were estimated from the community’s fluorescence measurements at 20 h for different concentrations of only one inducer added to the culture. Non-linear least squares optimization was used to fit a Hill equation to the dose response data from each input-output combination, and the optimized ℎ_i,j_and 𝐾_i,j_ values were saved.

Carrying capacity (𝑁_m_) was fixed to a value of 1.5 and the remaining parameters were estimated using non-linear least squares optimization to fit the full ODE model to all time course data. Initial estimates and bounds provided to the optimizer are in Supplementary Table 2.

### Simulated data generation

Using the optimized parameter set for each microbial sensor community, we simulated 10,000 OD and fluorescence time courses using solve_ivp function (SciPy). Chemical input concentrations were randomized on a uniformly distributed log scale within a range bounded by the lower 1 percentile and upper 99th percentile of the sensor’s dynamic range, as represented by the inequality,

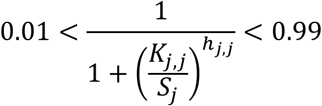

and normalized by the half maximum concentration, 𝐾_j,j_, that corresponds to the intended inducer-fluorescence pair. Initial conditions for OD and fluorescence were randomized on a normal distribution with mean, standard deviation, minimum and maximum limits set by the respective measured experimental values. Each simulation had the same number of time points as the corresponding experimental dataset. The OD and fluorescence time courses were concatenated to create a 1-dimensional vector per simulation that is of length 𝑡 x (𝑁 + 1), where 𝑁 is the number of fluorescence colors in the community and 𝑡 is the number of time points simulated.

### Secondary data processing

The experimental data was filtered to remove samples where the inducer concentration was equal to 0 or where the inducer concentration was below the 1 percentile or above the 99^th^ percentile of the sensor’s dynamic range as determined by the same inequality as above. The chemical input concentrations were normalized by the half maximum concentration, 𝐾_j,j_, that corresponds to the intended inducer-fluorescence pair. The filtered experimental data was combined with the simulated data to create the augmented dataset. With each time point of the time course data treated as a feature of the dataset, each feature was min-max scaled to lie within a range between 0 and 1. The chemical input concentration data was log_10_ transformed. Data was randomly shuffled with an 80/20 train-test split.

### VAE construction and training

The VAE utilizes a one-dimensional convolutional neural network (CNN) to encode and decode the time course data. The encoder consists of four convolutional layers with ReLU activation functions, progressively reducing the time course dimensionality to a 6-channel latent representation. Fully connected layers then compute the mean and log variance, mapping the representation to a 10-dimensional latent space. The reparameterization trick is applied to sample from the latent distribution for reconstruction. The decoder reconstructs the time courses from the latent space using a fully connected layer followed by transposed convolutional layers that upsample the latent representation back to the original time course dimensions. The model was trained with a batch size of 32 for 400 epochs using the Adam optimizer (learning rate = 1e-3, weight decay = 1e-5). The objective function combines reconstruction loss (MSE) and KL divergence to balance accurate reconstruction with regularization of the latent space. A warmup learning rate schedule was applied for the first 8 epochs, followed by exponential decay (decay rate, γ = 0.99) with a minimum learning rate of 5e-6. Early stopping was employed, where training was terminated if the test loss did not improve for 30 consecutive epochs. After training, latent representations were extracted by computing the mean vectors of the encoder output for both train and test datasets.

### Combined VAE-MLP construction and training

The MLP was trained to predict analyte concentrations from the latent representations that were output from the VAE. The MLP consisted of three fully connected linear layers. The first layer mapped the 10-dimensional latent space to a 128-dimensional hidden representation, followed by a ReLU activation function. The second layer of the same width further processed the hidden representation without reducing the dimensionality, followed by another ReLU activation. The final output layer mapped the hidden representation to the predicted analyte concentrations (2 or 3 dimensions). The MLP training was done after the VAE was already trained and frozen, to form a combined model, where the VAE’s resulting latent variables were passed through the MLP to generate analyte concentration predictions. The training procedure for the MLP, including the optimizer and learning rate scheduler, was identical to that of the VAE.

## Author Contributions

KED and LY conceptualized and designed the research and wrote the manuscript. KED conducted experimental data generation, computational pipeline development and analysis. ARS conducted experimental data generation and analysis. IS performed computational pipeline analysis and gave writing feedback. AR and RD conducted experimental data generation, aided with computational pipeline optimization, and gave writing feedback. HRM generated the antibiotic treatment data. KED, HVL, and SL cloned sensor circuits. HVL conducted preliminary data generation and helped conceptualize the research. SL and NG gave writing feedback. NG provided sink samples and protocols. DA and CG provided sink samples and protocols and provided guidance on applications and future directions. LY aided in computational pipeline development and analysis and experiment design.

## Acknowledgements

This work was partially supported by the National Institutes of Health (R01GM098642, R01EB031869, PI: You), Center for Disease Control (Cooperative Agreement No. U54CK000616, PI. Anderson), and the National Science Foundation (Cooperative Agreement No. EEC-2133504, PI: Gunsch). The funders had no role in study design, data collection and analysis, decision to publish, or preparation of the manuscript.

## Data availability

All data is available in the main text, supplementary information, or in GitHub at: https://github.com/ked46/multiplexed_sensing

## Code availability

Code used for the computational pipeline described here and for figure generation is available in GitHub at: https://github.com/ked46/multiplexed_sensing

