## Supplementary Information for "Engineering microbial consortia for distributed signal processing"

**Supplementary Materials for this manuscript include the following:**

Supplementary Note

Supplementary Tables 1-2

Supplementary Figures 1-14

**Supplementary Note**

VAE architecture and latent dimension optimization

To determine the optimal VAE architecture, we iteratively defined, trained, and tested the VAE with different architectures on the same dataset. Specifically, we tested architectures that varied in number of hidden layers and in the width of each hidden layer. We did this optimization for both the aTc-GFP and IPTG-mCherry community and the THS-CFP and TTR-YFP community datasets separately. Each architecture was trained in triplicate, and the resulting weights were saved. Architectures that yielded $R^2 > 0.989$ for reconstructions of the augmented training and testing time courses for that training iteration were subsequently used in the VAE-MLP framework. The latent dimensions from the architectures passing this threshold were used as inputs to train the MLP. The VAE architecture that yielded the highest average $R^2$ for predicting the experimental test set concentrations for both datasets and that passed the VAE reconstruction threshold more than once in both analyses was selected for the final VAE-MLP pipeline used in this study.

Using the optimized VAE architecture, we systematically tested VAE latent space sizes ranging from 5 to 20 dimensions to determine the optimal number for subsequent MLP predictions of input concentrations. We performed this optimization on both the aTc-GFP and IPTG-mCherry community and the THS-CFP and TTR-YFP community datasets. The VAE was trained three times for each latent space size, and the model was saved after each iteration. Latent variables that yielded $R^2 > 0.97$ for VAE reconstructions of the augmented training and testing time courses for that training iteration were subsequently used in the MLP to predict inputs. The latent dimension size that yielded the highest average $R^2$ for predicting experimental test set concentrations for both datasets was selected for the final pipeline used in this study.

### Supplementary Tables

#### Supplementary Table 1: Addgene references for all plasmids purchased for this study

| Plasmid | Addgene reference |
|---|---|
| pKD236-4b | Addgene plasmid # 90956; http://n2t.net/addgene:90956; RRID:Addgene_90956 |
| pKD237-3a-2 | Addgene plasmid # 90957; http://n2t.net/addgene:90957; RRID:Addgene_90957 |
| pKD238-1a | Addgene plasmid # 90958; http://n2t.net/addgene:90958; RRID:Addgene_90958 |
| pKD239-1g-2 | Addgene plasmid # 90959; http://n2t.net/addgene:90959; RRID:Addgene_90959 |
| pAJM.1642 | Addgene plasmid # 108535; http://n2t.net/addgene:108535; RRID:Addgene_108535 |
| pAJM.657 | Addgene plasmid # 108525; http://n2t.net/addgene:108525; RRID:Addgene_108525 |
| pAJM.011 | Addgene plasmid # 108529; http://n2t.net/addgene:108529; RRID:Addgene_108529 |
| pAJM.773 | Addgene plasmid # 108527; http://n2t.net/addgene:108527; RRID:Addgene_108527 |
| pAJM.847 | Addgene plasmid # 108524; http://n2t.net/addgene:108524; RRID:Addgene_108524 |
| pAJM.661 | Addgene plasmid # 108532; http://n2t.net/addgene:108532; RRID:Addgene_108532 |

#### Supplementary Table 2: Mechanistic model parameter values for initial estimate and bounds of the optimization

| Parameter | Initial estimate | Bounds |
|---|---|---|
| $\mu_i$ | 0.756 | 0.1 - 2 |
| $K_{s,i}$ | .099 | 0 - 1 |
| $\theta_i$ | 3 | 0 – 1 |
| $d_{S,j}$ | 0.5 | 0.0001 - 1 |
| $r_{0,i}$ | 0 | 0 - 1 |
| $k_i$ | 1 | 0.01 - 1 |
| $d_{p,i}$ | 0.1 | 0.6 - 6 |

### Supplementary Figures

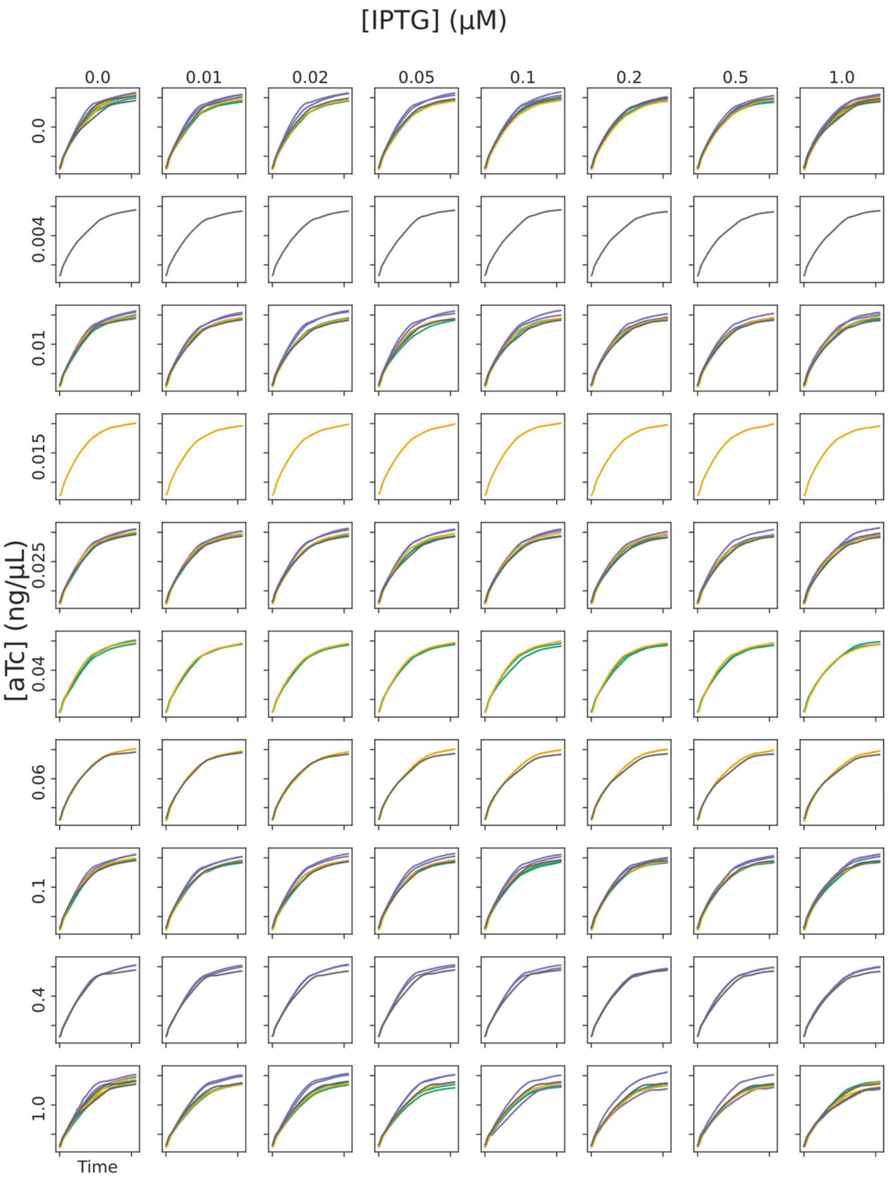


**Supplementary Figure 1.** OD time courses after data processing for all experimental conditions of the low-crosstalk aTc-GFP and IPTG-mCherry sensor set. Some samples here were excluded from full ODE model fitting and the VAE-MLP pipeline after determining the minimum and maximum threshold limits for the input concentrations based on the dose response curves (Methods).

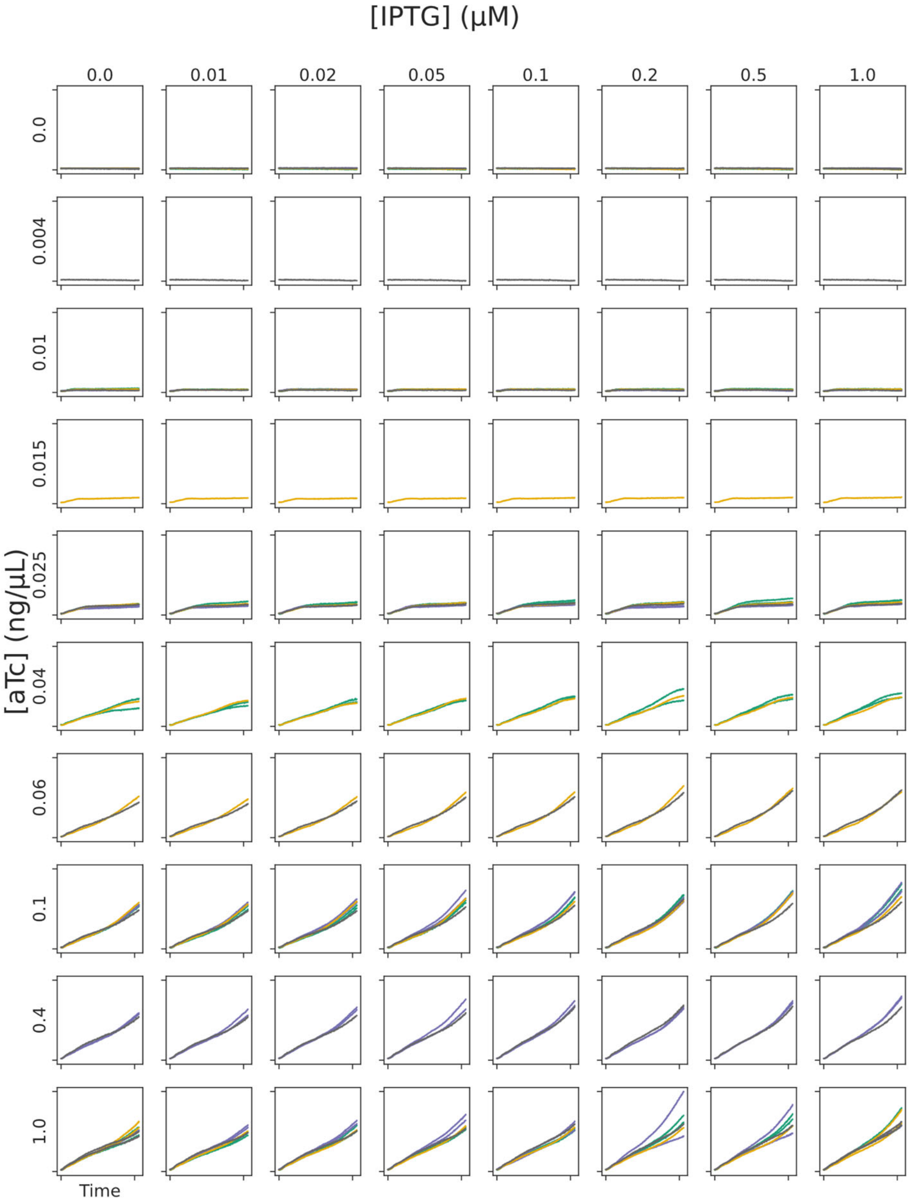


**Supplementary Figure 2.** GFP time courses after data processing for all experimental conditions of the low-crosstalk aTc-GFP and IPTG-mCherry sensor set. All samples here were used to estimate crosstalk and Hill equation parameters, but some were excluded from full ODE model fitting and the VAE-MLP pipeline after determining the minimum and maximum threshold limits for the input concentrations based on the dose response curves (Methods).

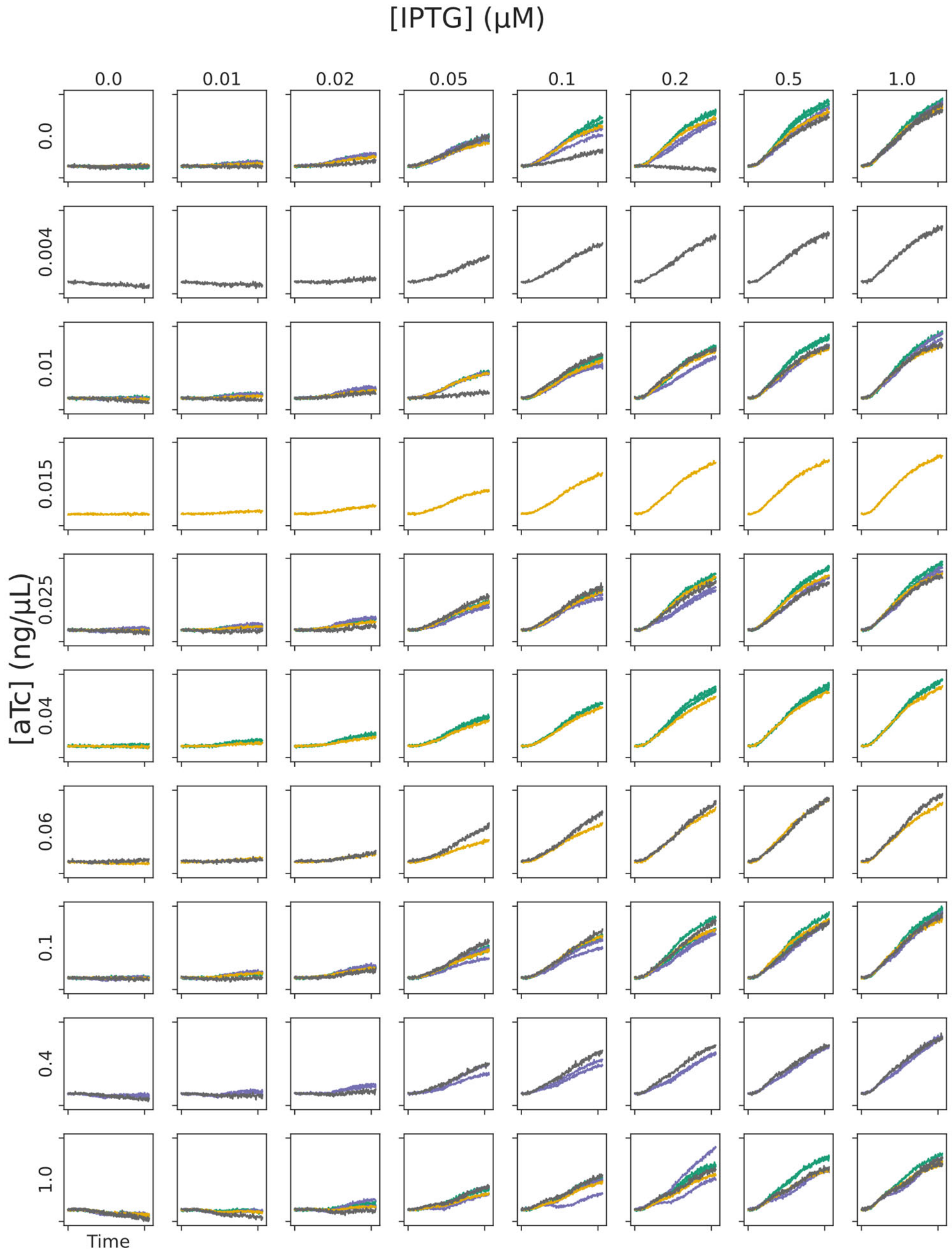


**Supplementary Figure 3.** mCherry time courses after data processing for all experimental conditions of the low-crosstalk aTc-GFP and IPTG-mCherry sensor set. All samples here were used to estimate crosstalk and Hill equation parameters, but some were excluded from full ODE model fitting and the VAE-MLP pipeline after determining the minimum and maximum threshold limits for the input concentrations based on the dose response curves (Methods).

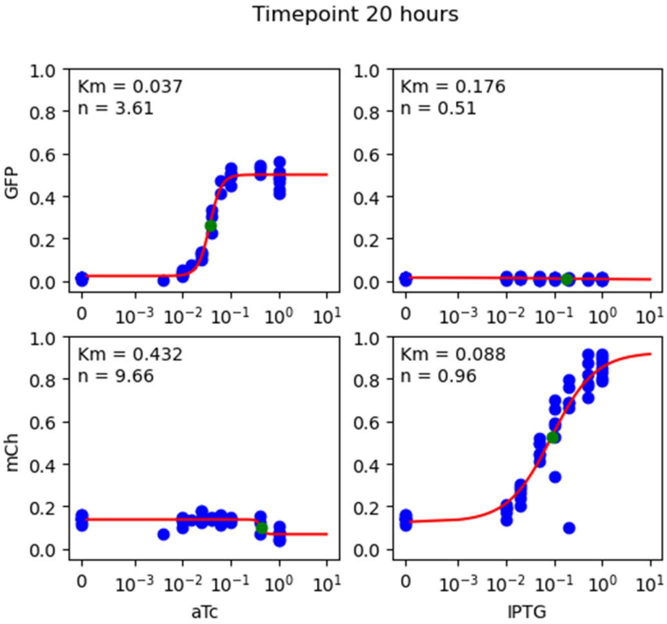


**Supplementary Figure 4.** Dose response data at 20 hours for aTc-GFP and IPTG-mCherry dataset. Blue points are experimental data, red line is the hill equation with parameters optimize to fit the experimental data. Km and n parameter values of the hill equation are estimated using this optimization.

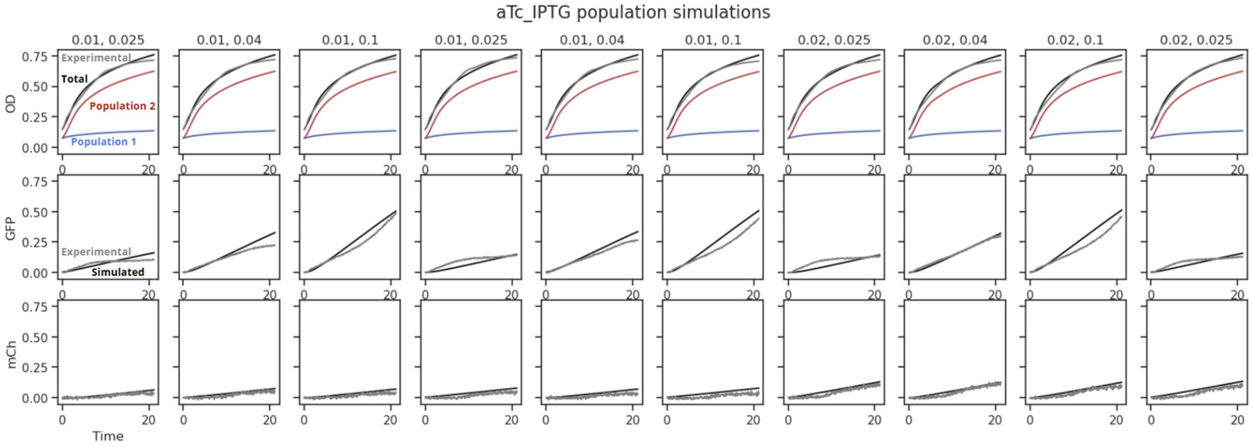


**Supplementary Figure 5.** Simulated growth curves of individual populations in the orthogonal microbial community for sensing aTc and IPTG, with different input concentrations. Population 1 (blue line) doesn't reach the same cell density as population 2 (red line). Total simulated OD (black line) and experimentally measured OD (gray line) are also plotted (top row). Simulated and experimental fluorescence curves are also plotted (middle and bottom row).

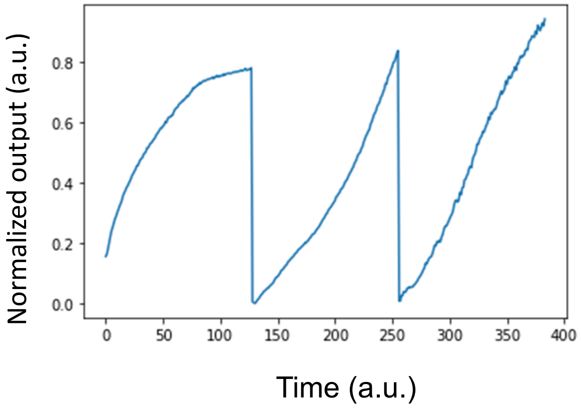


**Supplementary Figure 6.** Example of concatenated OD, GFP, and mCherry time courses, which form a one-dimensional data series.

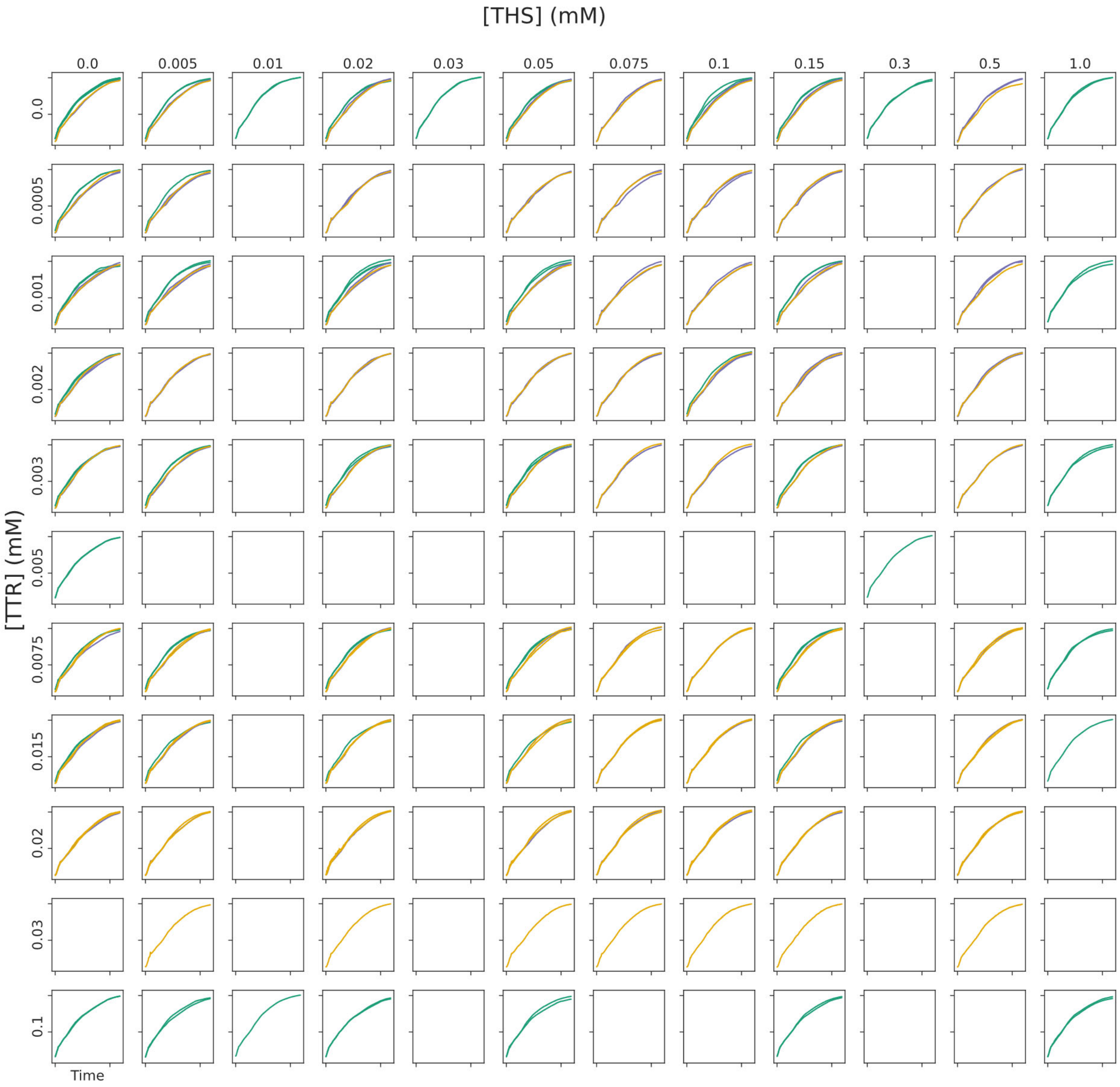


**Supplementary Figure 7.** OD time courses for all conditions from the high crosstalk ttr-ths sensor set after data processing, which were used as inputs into the VAE-MLP as well as for fitting the mechanistic model parameters to.

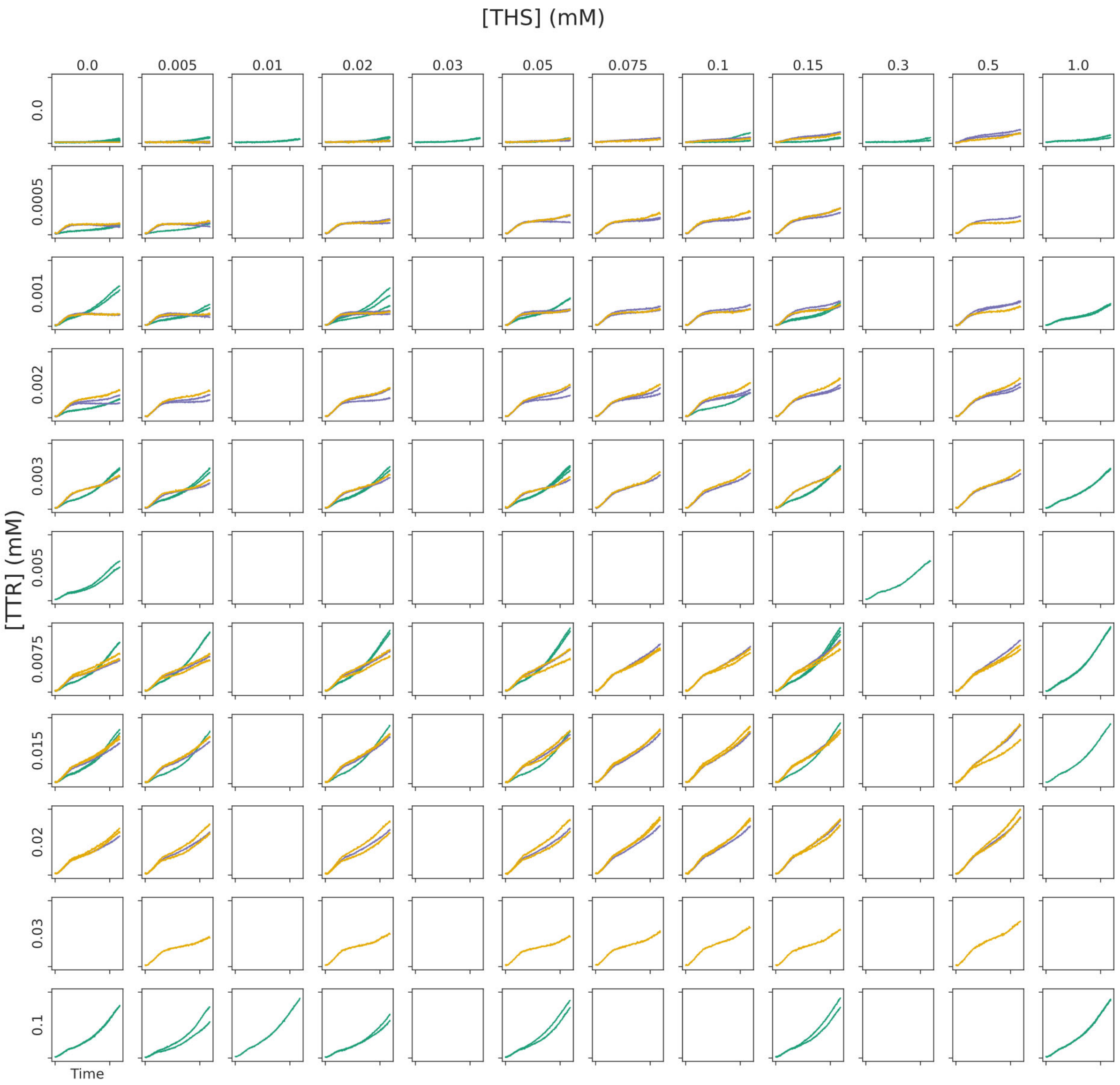


**Supplementary Figure 8.** YFP time courses for all conditions from the ttr-ths sensor set after data processing, which were used as inputs into the VAE-MLP as well as for fitting the mechanistic model parameters to.

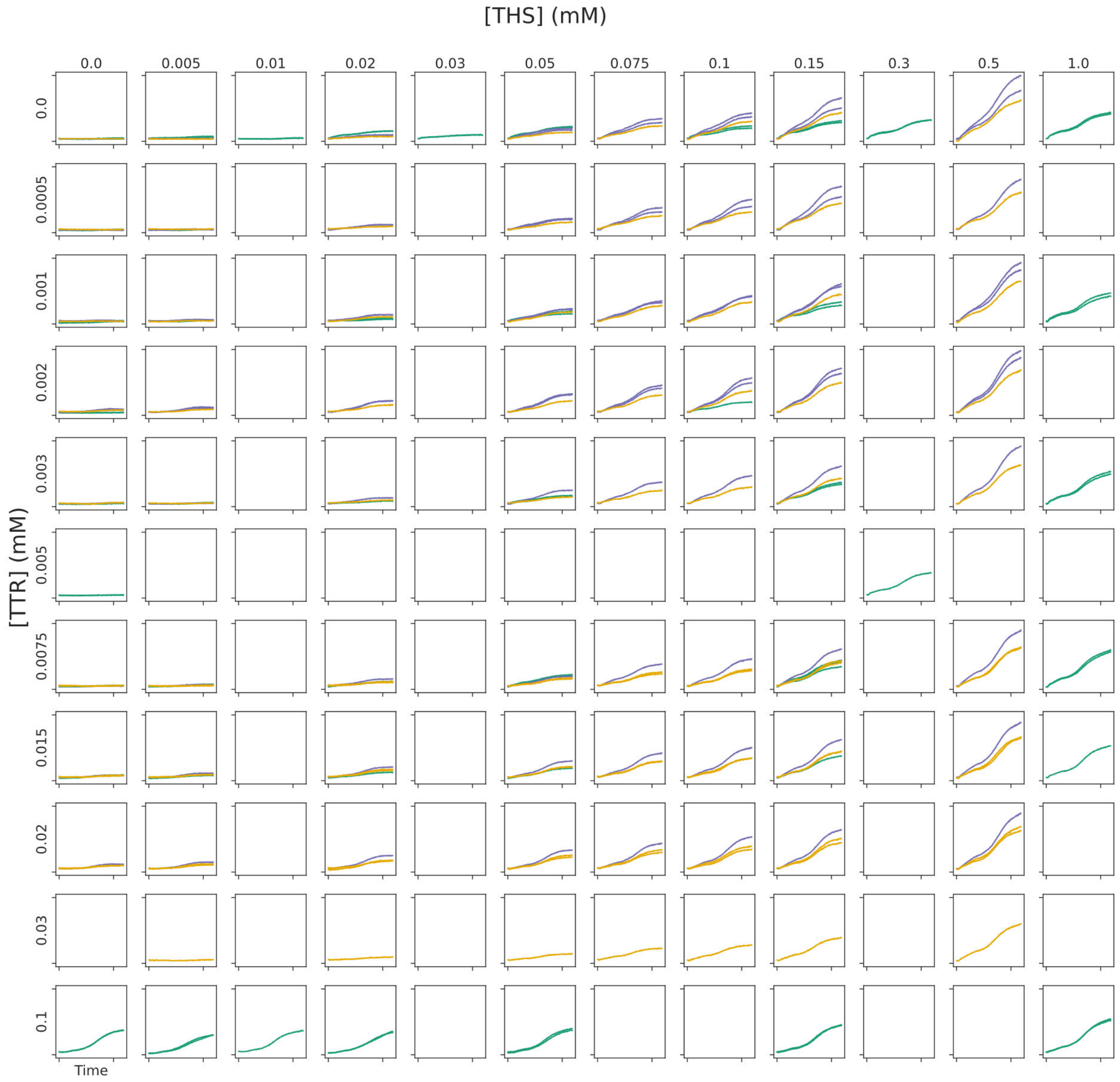


**Supplementary Figure 9.** CFP time courses for all conditions from the ttr-ths sensor set after data processing, which were used as inputs into the VAE-MLP as well as for fitting the mechanistic model parameters to.

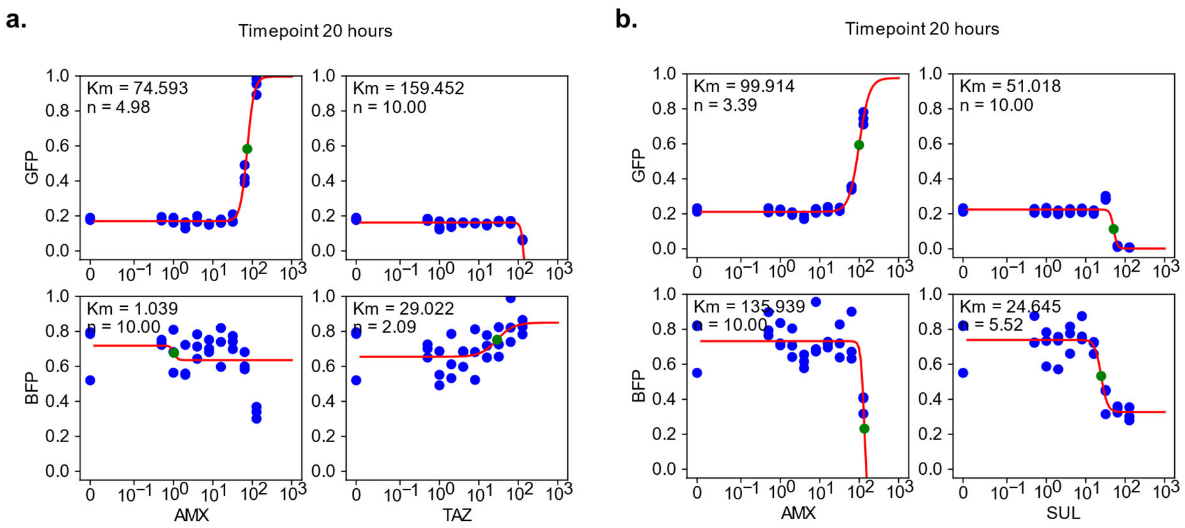


**Supplementary Figure 10:** Fluorescence dose responses of the antibiotic-treated microbial community when either antibiotic or inhibitor was added. Color bar and labeled value represent the fluorescence fold-change with respect to the fluorescence when neither antibiotic or inhibitor were added.

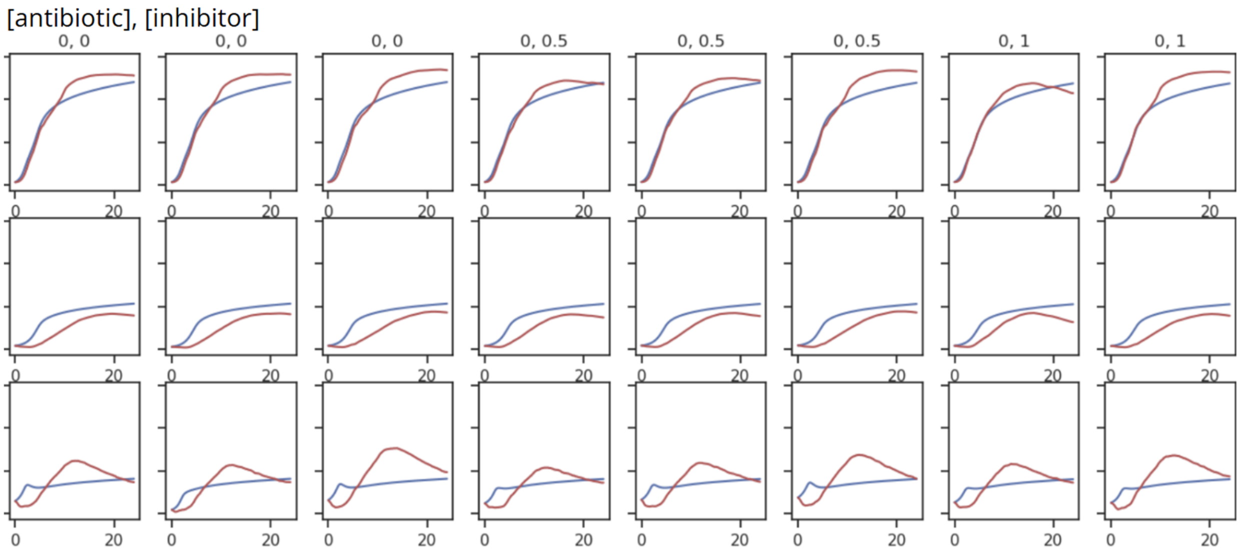


**Supplementary Figure 11:** Sampling of time course fitting results for antibiotic treatment data. Blue line is simulated data and red line is experimental data. OD (top row), GFP (middle row), and BFP (bottom row) are plotted.

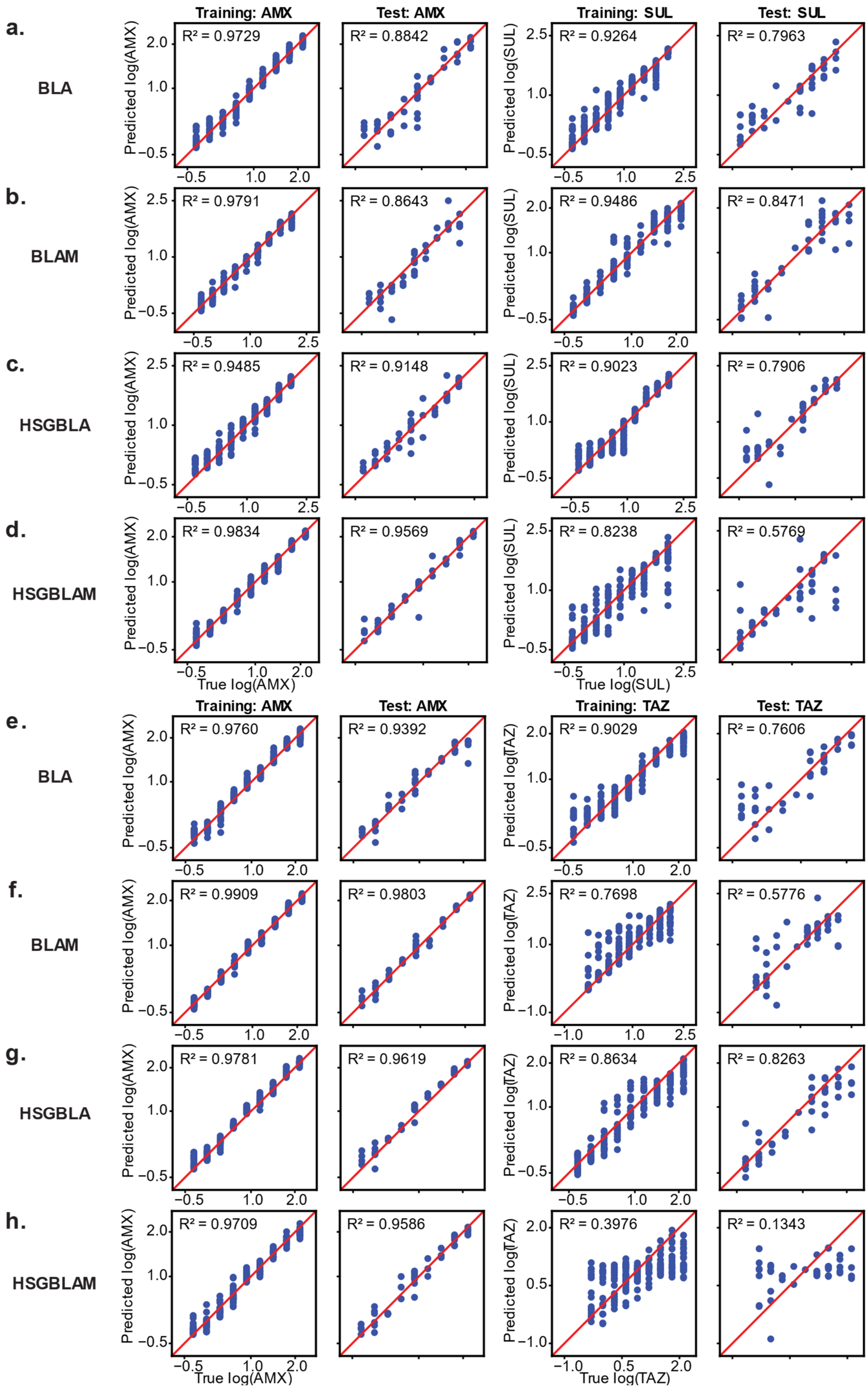
a.
BLA
b.
BLAM
c.
HSGBLA
d.
HSGBLAM
e.
BLA
f.
BLAM
g.
HSGBLA
h.
HSGBLAM
Training: AMX
Test: AMX
Training: SUL
Test: SUL
Training: TAZ
Test: TAZ
R² = 0.9729
R² = 0.8842
R² = 0.9264
R² = 0.7963
R² = 0.9791
R² = 0.8643
R² = 0.9486
R² = 0.8471
R² = 0.9485
R² = 0.9148
R² = 0.9023
R² = 0.7906
R² = 0.9834
R² = 0.9569
R² = 0.8238
R² = 0.5769
R² = 0.9760
R² = 0.9392
R² = 0.9029
R² = 0.7606
R² = 0.9909
R² = 0.9803
R² = 0.7698
R² = 0.5776
R² = 0.9781
R² = 0.9619
R² = 0.8634
R² = 0.8263
R² = 0.9709
R² = 0.9586
R² = 0.3976
R² = 0.1343
Predicted log(AMX)
Predicted log(SUL)
Predicted log(TAZ)
True log(AMX)
True log(SUL)
True log(TAZ)

**Supplementary Figure 12: VAE-MLP results of experimental antibiotic and inhibitor concentration predictions for all antibiotic/inhibitor datasets. a-d.** Predictions of the amoxicillin and sulbactam concentrations that were used to treat communities with different plasmids conferring antibiotic resistance in the resistant strain. **e-f.** Predictions of the amoxicillin and tazobactam concentrations that were used to treat communities with different plasmids conferring antibiotic resistance in the resistant strain. Resistance plasmids were Bla (**a,e**), BlaM (**b,f**), HSGBla (**c,g**), and HSGBlaM (**d,h**).

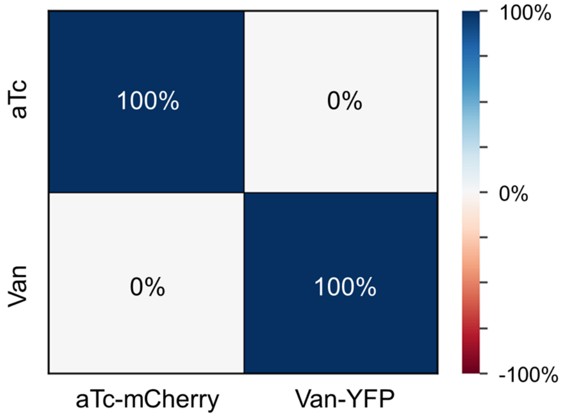


**Supplementary Figure 13:** Crosstalk heatmap for aTc-mCherry and Van-YFP sensor community responses to added Van and aTc in hospital sink wastewater. The two sensors do not have crosstalk with each other, but the complex background environment of the sink wastewater can introduce crosstalk.

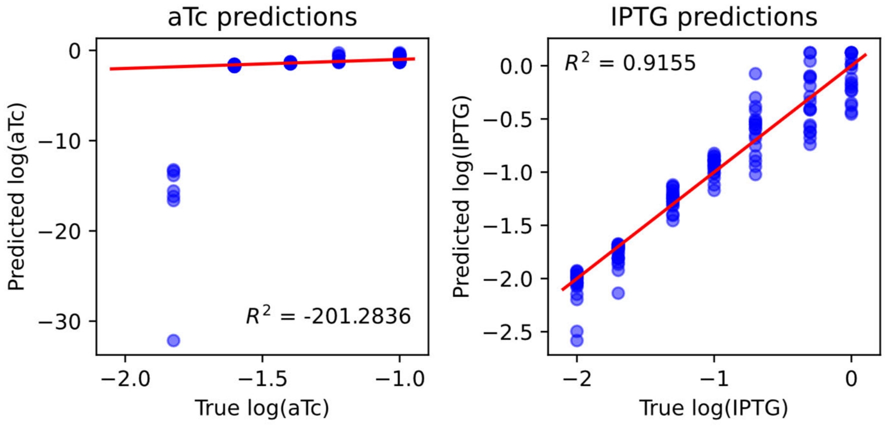


**Supplementary Figure 14: IPTG and aTc concentration predictions using non-linear optimization with the mechanistic model.** Using the optimized mechanistic model parameters for the IPTG-mCherry and aTc-GFP experimental dataset, we used a least squares optimization to estimate the IPTG and aTc concentrations of each sample. To fit the equations to the experimental curves, we only optimized IPTG and aTc concentrations to minimize residuals, keeping the parameters fixed. Initial guesses and bounds for both chemicals were the K value of the Hill equation and [0, 15] respectively. The aTc predictions were significantly lower fidelity than the predictions from the MLP, particularly for the lowest concentration. The mechanistic model also predicted IPTG concentrations with less accuracy than the MLP. $R^2$ values were calculated from the $\log_{10}$ transformed input values. Red line is Y=x for reference.
